# Mechanotransductive feedback control of endothelial cell motility and vascular morphogenesis

**DOI:** 10.1101/2022.06.15.496293

**Authors:** Devon E. Mason, Paula Camacho, Megan E. Goeckel, Brendan R. Tobin, Sebastián L. Vega, Pei-Hsun Wu, Dymonn Johnson, Su-Jin Heo, Denis Wirtz, Jason A. Burdick, Levi Wood, Brian Y. Chow, Amber N. Stratman, Joel D. Boerckel

## Abstract

Vascular morphogenesis requires persistent endothelial cell motility that is responsive to diverse and dynamic mechanical stimuli. Here, we interrogated the mechanotransductive feedback dynamics that govern endothelial cell motility and vascular morphogenesis. We show that the transcriptional regulators, YAP and TAZ, are activated by mechanical cues to transcriptionally limit cytoskeletal and focal adhesion maturation, forming a conserved mechanotransductive feedback loop that mediates human endothelial cell motility *in vitro* and zebrafish intersegmental vessel (ISV) morphogenesis *in vivo*. This feedback loop closes in 4 hours, achieving cytoskeletal equilibrium in 8 hours. Feedback loop inhibition arrested endothelial cell migration *in vitro* and ISV morphogenesis *in vivo*. Inhibitor washout at 3 hrs, prior to feedback loop closure, restored vessel growth, but washout at 8 hours, longer than the feedback timescale, did not, establishing lower and upper bounds for feedback kinetics *in vivo*. Mechanistically, YAP and TAZ induced transcriptional suppression of RhoA signaling to maintain dynamic cytoskeletal equilibria. Together, these data establish the mechanoresponsive dynamics of a transcriptional feedback loop necessary for persistent endothelial cell migration and vascular morphogenesis.

## Introduction

Christiane Nüsslein-Vohlard wrote in her book, Coming to Life, that *“during development, any change in cell shape is preceded by a change in gene activity”* (Nüsslein-Volhard, 2006). It is intuitive that gene expression is necessary to produce, for example, the cytoskeletal and adhesion machinery that enable cell motility, but whether persistent gene expression is required for continued migration is not. We previously found, to our surprise, that blockade of *de novo* gene expression (i.e., transcription or translation), caused progressive motility arrest (Mason et al., 2019). This requirement for continued transcription is executed by a feedback loop mediated by the transcriptional regulators, Yes-associated protein (YAP) and transcriptional-coactivator with PDZ-binding motif (TAZ) (Mason et al., 2019). Here, we define the kinetics of this mechanotransductive feedback loop, *in vitro* and *in vivo*.

Mechanotransduction is often evaluated after the cell has reached equilibrium and represented as a one-way path from cytoskeletal activation to nuclear information transfer, resulting in a state variable shift (e.g., lineage commitment). However, mechanotransduction can also activate feedback loops, which alter how the cell responds to subsequent mechanical stimuli. These feedback loops are necessary to maintain a responsive cytoskeletal equilibrium or to preserve an adapted state.

Recently, we and, independently, the Huveneers group found that human endothelial cells maintain cytoskeletal equilibrium for persistent motility through a YAP/TAZ-mediated feedback loop that maintains tensional homeostasis (Mason et al., 2019; van der Stoel et al., 2020). Because YAP and TAZ are activated by tension of the cytoskeleton (Dupont et al., 2011), suppression of cytoskeletal tension by YAP/TAZ transcriptional target genes constitutes a negative feedback loop that acts as a control system to maintain cytoskeletal homeostasis for persistent motility via modulation of Rho-ROCK-myosin II activity (Fig. 1A) (Mason et al., 2019). Both we and the Huveneers group found that perturbation of genes and pathways regulated by YAP/TAZ mechanotransduction can functionally rescue motility in YAP/TAZ-depleted cells (e.g., RhoA/ROCK/myosin II, NUAK2, DLC1) (Mason et al., 2019; van der Stoel et al., 2020). These prior findings establish functional nodes of this mechanotransductive feedback loop.

**Figure 1:**
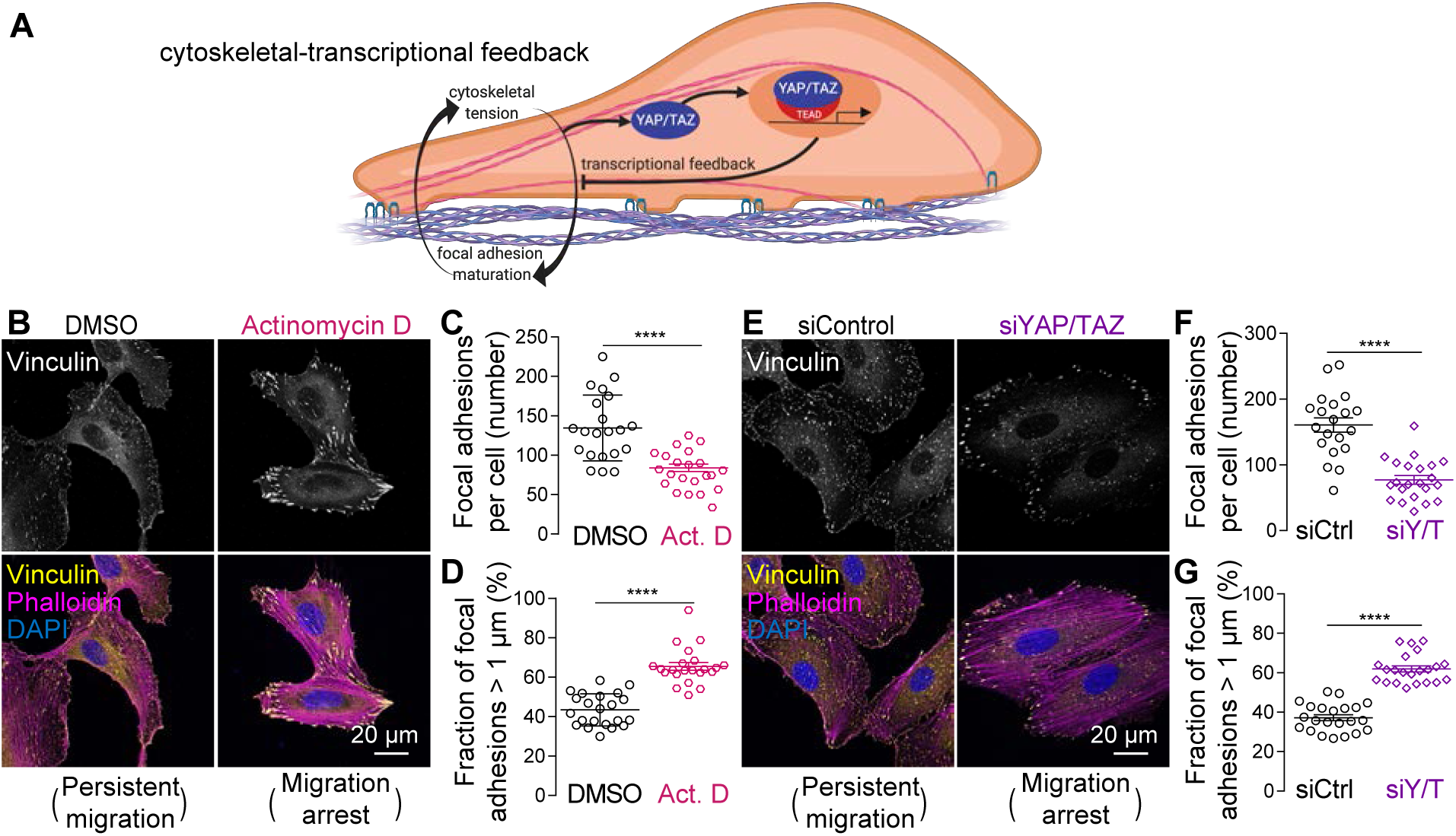
Cytoskeletal-transcriptional feedback prevents excessive focal adhesion maturation to enable persistent motility in human ECFCs. **A.** Model of YAP/TAZ-mediated transcriptional feedback regulation of the cytoskeleton, as established in (Mason et al., 2019; van der Stoel et al., 2020). **B-D.** ECFCs were treated with the transcription inhibitor, Actinomycin D, and stained for vinculin, phalloidin (F-actin), and DAPI (nuclei) after 24 hours. **E-G.** ECFCs were depleted of both YAP and TAZ by RNAi and stained for vinculin, phalloidin (F-actin), and DAPI (nuclei) after 28 hours. Both transcription inhibition and YAP/TAZ depletion induced motile arrest, indicated by loss of fan-shaped lamellipodial migration and acquisition of an oval shape, ringed by mature focal adhesions (**B, E**). Vinculin+ focal adhesions (**C, F**) and mature vinculin+ focal adhesions, defined as greater than 1 µm (**D, G**). N = 20-22 cells per condition. **** p < 0.0001, Student’s two-tailed unpaired t-test. Data are shown as mean ± S.E.M.

Here, we sought to determine the characteristic time scales and physiologic impacts, using two orthogonal model systems: human endothelial colony forming cells *in vitro* and zebrafish intersegmental vessel development *in vivo.* Endothelial colony forming cells (ECFCs) are circulating endothelial cells that activate in response to vascular injury, adhering and migrating to the injury site to repopulate the damaged endothelium (Ingram et al., 2004; David A Ingram et al., 2005). To study the dynamics of cytoskeletal feedback control in ECFCs, we performed dynamic quantification of cytoskeletal and cellular morphodynamics over time after adhesion. To study the dynamics of vascular morphogenesis, we performed longitudinal, quantitative imaging of intersegmental vessel (ISV) development. This model is particularly tractable for this question as zebrafish ISV development exhibits similar migratory kinetics to human ECFC migration *in vitro* and can be quantified longitudinally in fluorophore-labeled endothelial cells *in vivo* (Phng et al., 2013; Rosa et al., 2022).

Here, we define the characteristic time scales of YAP/TAZ-mediated transcriptional feedback loop closure and demonstrate conservation of feedback loop mediators and kinetics between human endothelial cell migration *in vitro* and zebrafish vascular morphogenesis *in vivo*.

**Movie 1: Representative effects of transcription inhibition on ECFC migration and cytoskeletal dynamics.** Adherent ECFCs, transfected with LifeAct-tdTomato, were treated with DMSO (left) or the transcription inhibitor, Actinomycin D (right). Images were taken in 3-minute intervals for 12.5 hours after inhibitor treatment. Time is shown in the top left as hours:minutes.

### Transcriptional feedback control of cytoskeletal and focal adhesion maturation

To test how *de novo* gene expression mediates endothelial cell motility, we evaluated live migration of human ECFCs for 10 hours after broad transcription inhibition by Actinomycin D. Control cells exhibited dynamic actin stress fiber formation and remodeling (via LifeAct-tdTomato) and persistent motility, while transcription-inhibited cells initiated with similar dynamics, but exhibited stress fiber stabilization and motile arrest between four and eight hours after transcription blockade (Movie 1). Next, we evaluated the effects of transcription-inhibited stress fiber arrest on the formation and maturation of focal adhesions at twenty-four hours after *in situ* Actinomycin D treatment (Fig. 1B-D). *In situ* transcription inhibition decreased the total number of vinculin+ adhesion plaques (Fig. 1C) and increased the fraction of adhesions over 1 µm in length, a measure of focal adhesion maturity (Fig. 1D).

As with broad transcription inhibition, we previously found that *in situ* depletion of YAP and TAZ by RNAi arrested cell motility, illustrated here by live-migration sparklines over 10 hours: siControl: 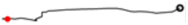, siYAP/TAZ: 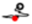(25 μm scale-bar: –). (Mason et al., 2019) *In situ* YAP/TAZ depletion decreased the total number of vinculin+ adhesion plaques, and increased the fraction of mature adhesions (Fig. 1E-G). Control cells exhibited fan-shaped lamellipodial migration while both transcription inhibition and YAP/TAZ depletion impaired polarization, and induced robust ventral stress fiber formation and peripheral focal adhesion maturation. Together with our prior findings (Mason et al., 2019), these data demonstrate that continued gene expression, via YAP/TAZ-mediated transcription, maintains cytoskeletal equilibrium for persistent motility.

**Movie 2: Representative cytoskeletal dynamics of LifeAct-tdTomadto-expressing ECFCs during first hour after adhesion to glass cover slip.** Images were taken in 10-second intervals 1 hour after attachment. Time is shown as minutes:seconds.

### Mechanoregulation of endothelial cell morphodynamics and persistent motility

To study the dynamics of cytoskeletal feedback control in ECFCs, we developed an adhesion-spreading-polarization-migration (ASPM) assay (Fig. 2A), following (Benecke et al., 1978). This assay corresponds to ECFC adhesive function *in vivo* and features cell trypsinization and reattachment at t_0_ to re-activate the cytoskeleton from a partially reset state. Upon adhesion, ECFCs initially spread isotropically, then polarize and transition to randomly-directed motility (Movie 2).

**Figure 2:**
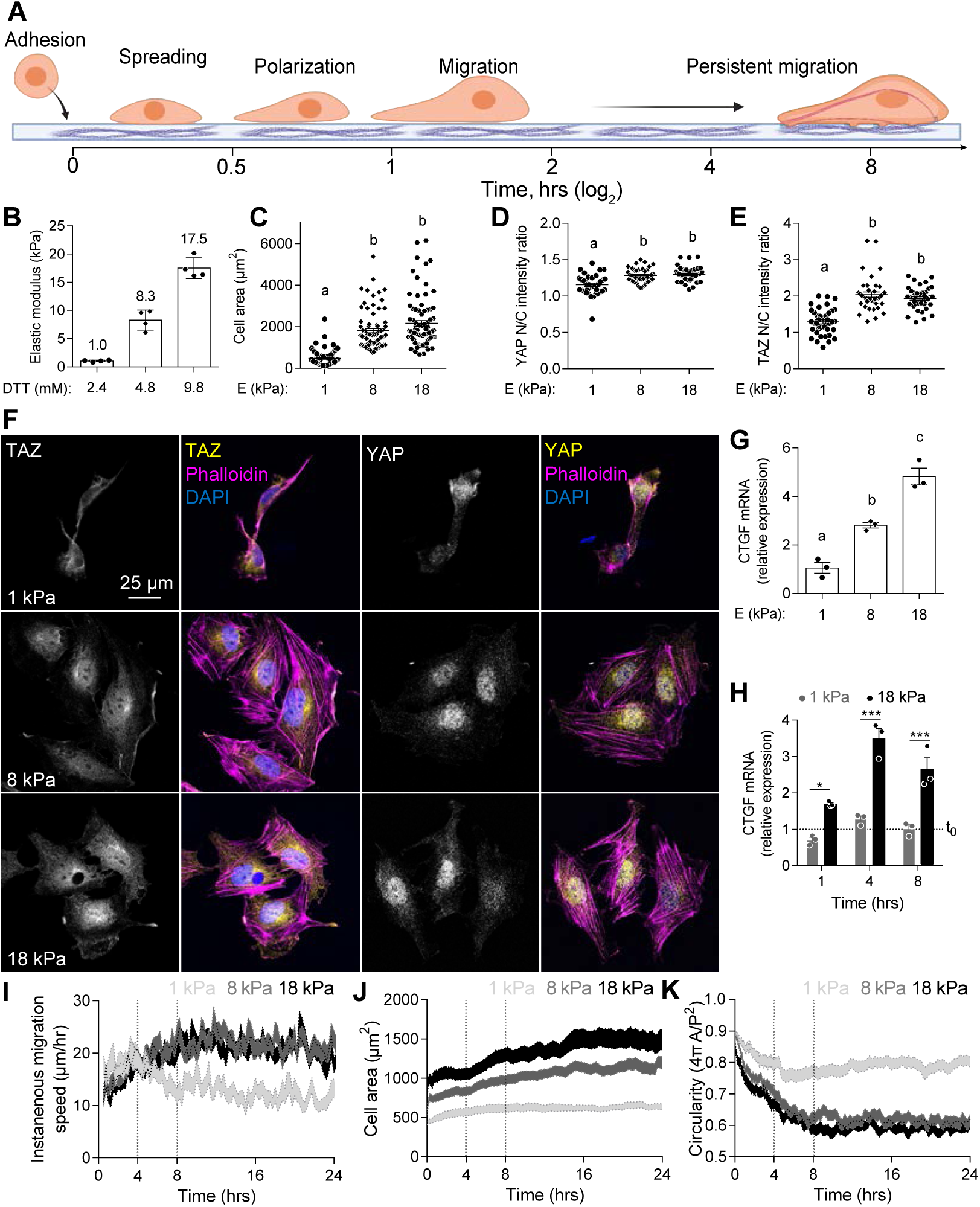
Extracellular matrix rigidity regulates mechanotransduction and motility dynamics in human ECFCs. **A.** Schematic of adhesion-spreading-polarization-migration (ASPM) assay. **B.** Methacrylated hyaluronic acid (MeHA) hydrogels were cross-linked with varying concentrations of DTT to form 1 kPa (soft), 8 kPa (moderate), and 18 kPa (stiff) hydrogels. Atomic force microscopy-measured elastic moduli are shown as mean ± S.D. n = 4, p < 0.0002, one-way ANOVA with Tukey’s post hoc test. ECFCs were then seeded on the MeHA hydrogels and assayed at 4 hours post-attachment. **C.** Cell area. n = 80 cells, p < 0.0001, Kruskal-Wallis with Dunn’s post hoc test. **D-E.** Quantification of the nuclear-to-cytosolic ratio of fluorescent intensity for YAP (D) and TAZ (E) at 4 hours post-attachment. n = 40 cells, p < 0.0001, one-way ANOVA with Tukey’s post hoc test. **F.** Representative immunofluorescent images visualizing F-actin (magenta), YAP or TAZ (yellow), and nuclei (blue) at 4 hours post-attachment. **G.** qPCR for the YAP/TAZ-target gene, CTGF. n = 3, p < 0.0054, one-way ANOVA with Tukey’s post hoc test. **H.** CTGF mRNA expression at 0, 1, 4, and 8 hours post-seeding, compared to unattached cells at t_0_ (dotted line). N = 3, * p < 0.01, *** p < 0.0008, two-way ANOVA with Sidak’s post hoc test. **I-K.** Next, motility of mTomato-expressing ECFCs was tracked over 20 hours post-attachment. Instantaneous migration speed, cell area, and circularity were calculated at 15-minute intervals until hour 24. Soft (n = 88 cells), moderate (n = 86 cells), and stiff (n = 89 cells). Data are shown as mean ± S.E.M. in error bars or shaded bands.

To determine how matrix mechanical cues regulate the kinetics of ECFC mechanotransduction, motility, and morphodynamics, we synthesized hyaluronic acid-based hydrogels with rigidities of 1, 8, and 18 kPa (soft, moderate, and stiff, respectively) (Fig. 2B). These stiffnesses were selected to correspond to the stiffness variation of the perivascular extracellular matrix from early development to adulthood (Huynh et al., 2011) and span the commonly-observed inflection point for mechanotransductive transition (Cosgrove et al., 2016). Compared to 1kPa, ECFCs seeded on 18 kPa matrices had increased cell area (Fig. 2C; p < 0.0001), YAP nuclear localization (Fig. 2D; p < 0.0001), TAZ nuclear localization (Fig. 2E; p < 0.0001), and mechanotransductive YAP/TAZ-TEAD-induced connective tissue growth factor (CTGF) mRNA gene expression (Fig. 2F; p < 0.0001). Cell morphology and YAP/TAZ nuclear localization were not statistically distinguishable between 8 and 18 kPa, though CTGF induction was significantly greater on 18 kPa matrices at 4 hours post-attachment (Fig. 2G; p < 0.0027).

To determine the dynamics of YAP/TAZ mechanotransduction by adhesion to varying-stiffness substrates, we trypsinized cultured ECFCs and adhered them at t_0_ to 1, 8, or 18 kPa MeHA hydrogels for 1, 4, or 8 hours. CTGF mRNA was elevated on 18 kPa vs 1 kPa hydrogels as early as 1 hour after attachment (p < 0.01), highest at 4 hours (p < 0.0001), and remained elevated at 8 hours after adhesion (p < 0.0001) (Fig. 2H). Compared to unattached t_0_ cells, CTGF expression was not significantly altered on 1 kPa substrates at any time (dotted line, p > 0.13).

Next, to determine how matrix mechanosensing impacts ECFC motility and morphodynamics, we tracked migration speed, spreading, and shape of mTomato-expressing ECFCs on 1, 8, or 18 kPa MeHA matrices from the time of attachment to 24 hours post-adhesion (Fig. 2I-K, Movies 3, 4). Regardless of matrix rigidity, cell motility and morphodynamics seemed to exhibit an inflection point at 4 hours post-attachment, and reache motile and morphological equilibrium by hour 8. To explore this equilibrium, we measured the difference in motility and morphology parameters between 8 hours and time zero (Δ_8-0_).

**Movies 3, 4: Effects of transcription inhibition on ECFC migration on soft, moderate, and stiff matrices.** mTomoto-expressing ECFCs were treated with DMSO (Movie 3) or the transcription inhibitor, Actinomycin D (Movie 4). Images were taken in 3-minute intervals for 12.5 hours after inhibitor treatment. Time is shown as hours:minutes.

Prior studies show that, at equilibrium, cells generally migrate faster on stiff substrates than on soft, and from softer toward stiffer matrices (Lange and Fabry, 2013; Sunyer et al., 2016; Sunyer and Trepat, 2020). However, ECFC migration speed was initially higher on soft substrates than on moderate or stiff substrates (average initial speed = 19.40, 9.00, and 8.93 µm/hr, for 1, 8, and 18 kPa respectively; Fig. 2I). Cell motility on soft hydrogels initially increased with time, but inflected at 4 hours and reached an equilibrium low by 8 hours. In contrast, cells on moderate and stiff hydrogels increased continuously, reaching an equilibrium high by 8 hours post-adhesion (Δ_8-0_ cell speed = -9.71, +12.92, and +12.39 µm/s for 1, 8, and 18 kPa respectively). Cell morphology dynamics exhibited similar kinetics with inflection points at 4 and 8 hours after attachment, reaching morphological equilibrium by hour 8 (Δ_8-0_ cell area = +197, +525, and +720 µm^2^; for 1, 8, and 18 kPa respectively; Fig. 2J). Similarly, cell circularity (inverse of elongation) decreased as a function of substrate rigidity, reaching morphological equilibrium by hour 8 (Δ_8-0_ circularity = -0.12, -0.26, and -0.24 for 1, 8, and 18 kPa respectively; Fig. 2K). These data implicate extracellular matrix mechanical cues as regulators of mechanotransductive, morphological, and motile adaptation.

### Transcriptional control of vascular morphogenesis

Mechanical stimuli and endothelial cell mechanotransduction are also critical for proper vascular morphogenesis *in vivo* (Boselli et al., 2015). Therefore, we next asked whether the YAP/TAZ-mediated cytoskeletal-transcriptional feedback loop identified in human EC migration *in vitro* is conserved *in vivo*. We measured dynamic vascular morphogenesis using embryonic zebrafish intersegmental vessel (ISV) migration using live imaging of genetically-encoded GFP-labeled endothelial cells (Tg*(fli:egfp) ^y1^*) (Fig. 3A) (Lawson and Weinstein, 2002). First, we exposed zebrafish embryos at 29 hours post-fertilization (hpf) to Actinomycin D and Puromycin to block general transcription and translation, respectively. Control embryos were treated with vehicle DMSO. Both Act. D (at 25 µg/ml (high), but not 10 µg/ml (low)) and Puromycin treatment slowed the kinetics of ISV growth by 35 hpf (Fig. 3C-E). Next, to inhibit YAP/TAZ signaling, we treated embryos with verteporfin (VP). Previously, we showed that VP treatment of ECFCs *in vitro* caused progressive cytoskeletal tension, focal adhesion maturation, and motility arrest equivalent to RNAi-mediated depletion (Mason et al., 2019), supporting use of this compound as an endothelial YAP/TAZ inhibitor. Exposure of zebrafish embryos to 50 µM VP at 29 hpf arrested vascular morphogenesis by 35 hpf (Fig. 3F-H). To block upstream YAP/TAZ activation by RhoA-ROCK-myosin signaling, we also treated with ROCK (Rockout) and non-muscle myosin II (Blebbistatin) inhibitors. Like VP, both ROCK and myosin inhibition abrogated ISV growth at 32 and 35 hpf (Fig. 3F-H). Together, these data confirm that the cytoskeletal-transcriptional feedback loop that mediates persistent human ECFC migration is conserved in zebrafish vascular morphogenesis.

**Figure 3:**
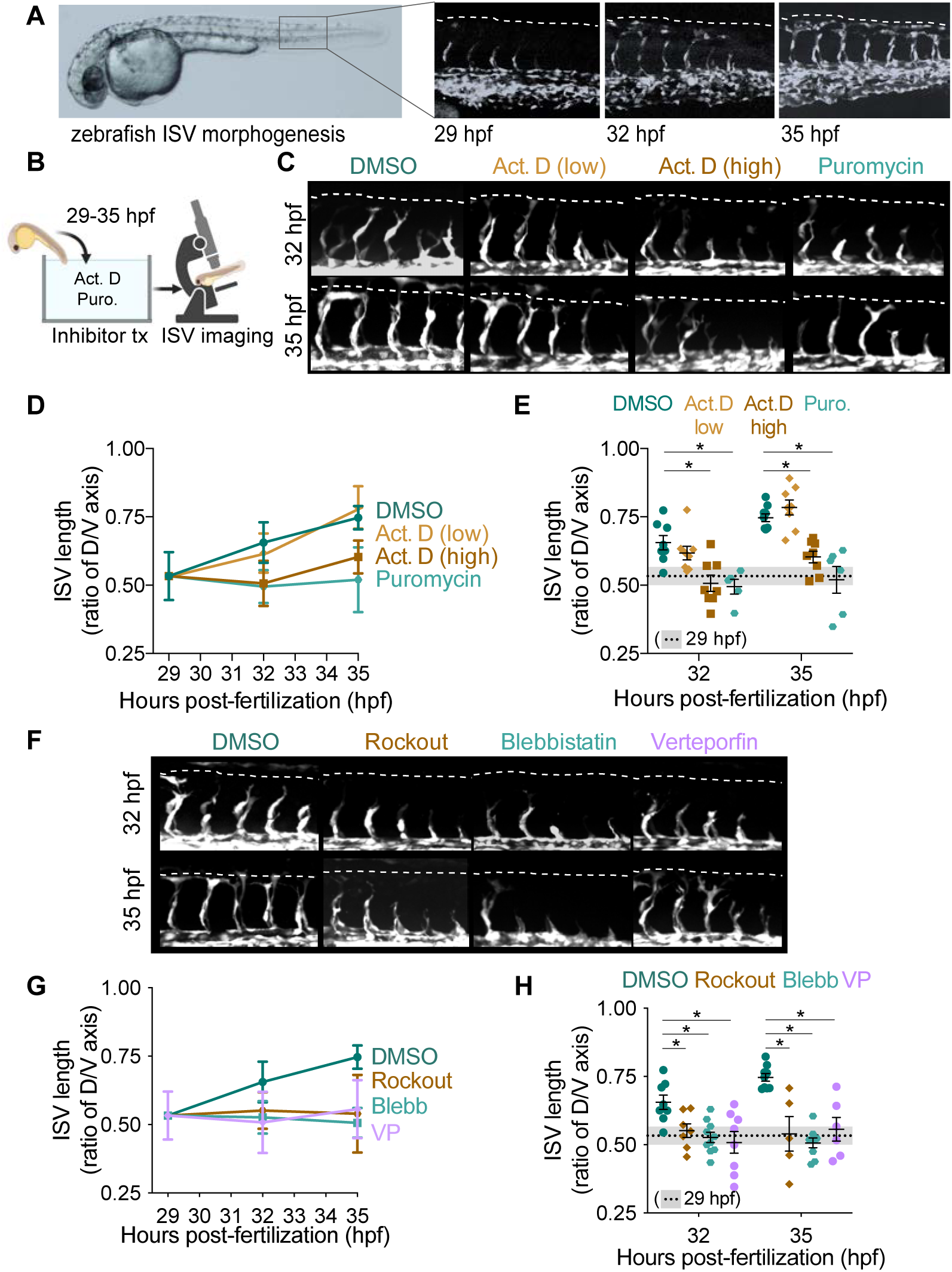
*De novo* gene expression and RhoA-YAP/TAZ signaling regulate zebrafish ISV morphogenesis *in vivo*. **A.** Bright field and spinning disc confocal imaging of the (Tg*(fli:egfp) ^y1^*) zebrafish transgenic line at 29, 32, and 35 hpf outlining the timeline of ISV morphogenesis in the zebrafish trunk. **B.** Schematic of experimental design. **C**. Representative spinning disc confocal images of DMSO, Actinomycin D (Act. D) 10 µg/ml (low), Act. D 25 µg/ml (high) and Puromycin treated zebrafish embryos. Compounds were added at 29 hpf and ISV morphogenesis imaged at 32 hpf (top) and 35 hpf (bottom). **D,E**. Quantification of ISV length, tracking individual fish over time, with data plotted as mean ± S.D. (D), vs comparative analysis at isolated time points with data plotted as mean ± S.E.M. * p < 0.05, two-way ANOVA with Sidak’s post hoc test (E). DMSO n = 8; Act. D (low) n = 8; Act. D (high) n = 8; and Puro n = 6. **F**. Representative spinning disc confocal images of DMSO, Rockout, Blebbistatin (Blebb), and Verteporfin (VP) treated zebrafish embryos. Compounds were added at 29 hpf and ISV morphogenesis imaged at 32 hpf (top) and 35 hpf (bottom). **G,H**. Quantification of ISV length, tracking individual fish over time, with data plotted as mean ± S.D. (G), vs comparative analysis at isolated time points with data plotted as mean ± S.E.M. * p < 0.05, two-way ANOVA with Sidak’s post hoc test (H). DMSO n = 8; Rockout n = 7; Blebb n = 10; and VP n = 8. Data points represent the ratio of ISV sprout length at the indicated time point to the total possible sprout length (dotted white line). Five ISVs were averaged per embryo to generate a single data point. ISV length ratio of ‘0’ indicates no ISV formation and ratio of ‘1’ indicates completion of ISV morphogenesis.

### Mechanotransductive feedback control of cytoskeletal and adhesion maturation regulates endothelial cell morphodynamics and persistent motility

Next, we tested how the mechanical environment influences the dynamics of feedback-inhibited motility arrest *in vitro*. To this end, we performed acute transcription inhibition at the time of cellular adhesion to MeHA hydrogels of 1, 8, or 18 kPa and tracked live cell migration, morphodynamics, and cytoskeletal and adhesion formation and maturation over 24 hours (Fig. 4A, Movies 3, 4). Matrix mechanotransduction responds with sigmoidal behavior to varied substrate rigidity (Cosgrove et al., 2016), and we observed similar dynamic responses to 8 and 18 kPa matrices, indicating these values both sit near or after the responsive plateau (Supplementary Fig. S1). For clarity and simplicity, we show responses to 1 and 18 kPa matrices (Fig. 4) and all groups in the supplement (Supplementary Fig. S1).

**Figure 4:**
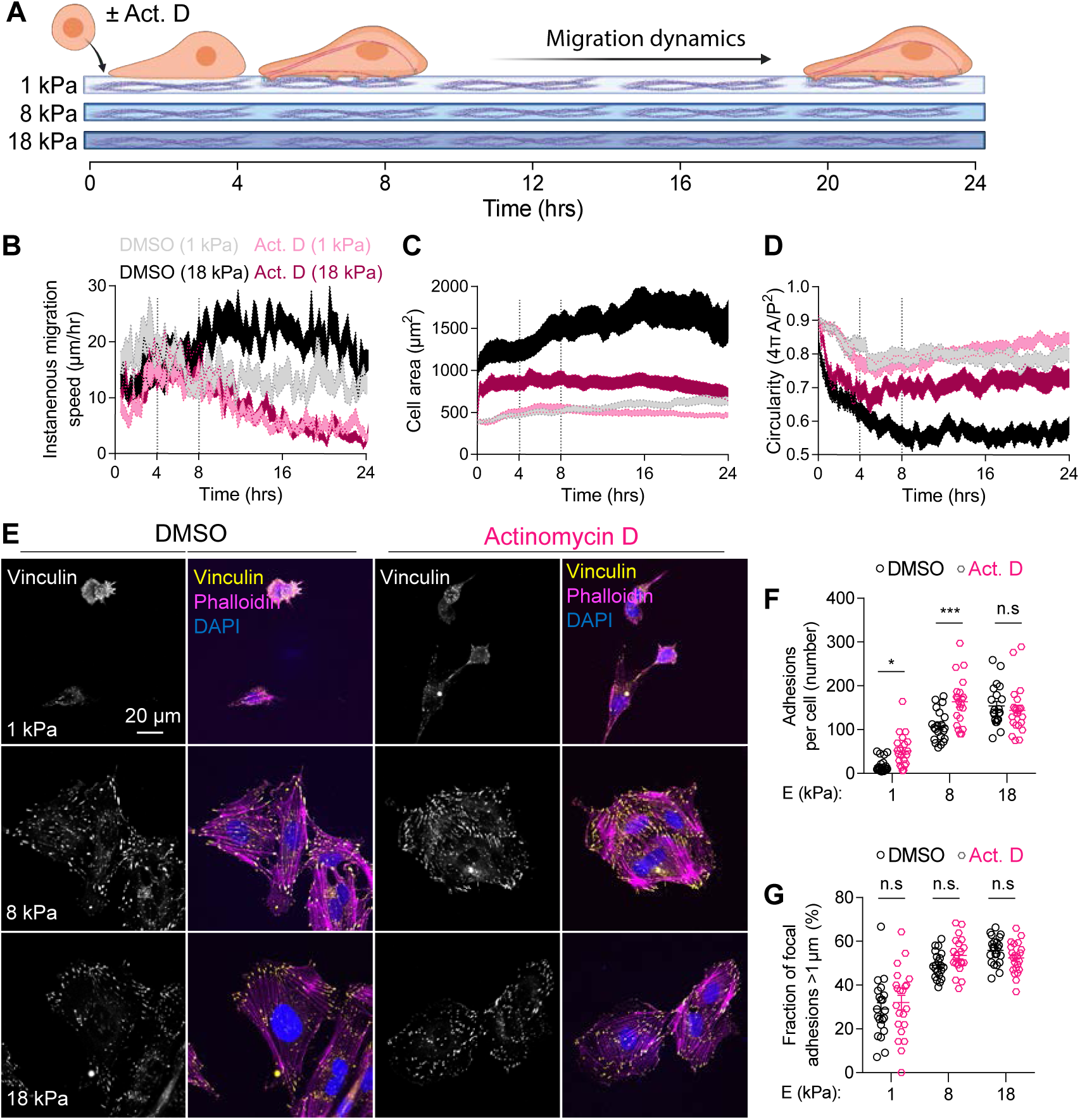
Acute transcription inhibition of human ECFCs induces motility and morphodynamic inflection at 4 hours post-attachment, followed by progressive motility arrest, on both soft and stiff substrates. **A.** Experiment schematic: mTomato-expressing ECFCs were treated with DMSO or Actinomycin D (Act. D; 0.1 µg/mL) at the time of seeding on 1, 8, or 18 kPa MeHA matrices. Cell morphology and migration were tracked for 24 hours post-attachment or fixed for immunofluorescent imaging at 4 hours post-attachment. **B-D.** Instantaneous migration speed, cell area, and circularity were calculated at 15-minute intervals until hour 24 in four groups: soft-DMSO (n = 48 cells), soft-Act.D (n = 63 cells), stiff-DMSO (n = 41 cells), and stiff-Act.D (n = 36 cells). Moderate stiffness groups are shown in Fig. S1. **E.** Immunofluorescent imaging of F-actin (magenta), vinculin (yellow), and nuclei (blue) at 4 hours post-attachment. **F, G.** Quantification of vinculin+ focal adhesions (F) and mature vinculin+ focal adhesions (G), defined as focal adhesions greater than 1 µm in length. n = 21-22, * p < 0.05, *** p < 0.0002, two-way ANOVA with Sidak’s post hoc test. Data are shown as mean ± S.E.M.

Cell motility and morphodynamics depended on both matrix rigidity and transcriptional feedback. Transcription inhibition reduced cell motility, spreading, and polarization, resulting in progressive motility arrest, regardless of matrix rigidity (Fig. 4B-D). On 18 kPa substrates, motility arrest initiated at 4 hours after adhesion, and by the time of motile equilibrium in control cells (8 hours after attachment), reduced migration speed relative to controls (Δ_8-0_ cell speed on 18 kPa = +5.44 and -9.20 µm/s for DMSO and Act. D, respectively). On 1 kPa substrates, transcription-inhibited cells similarly transitioned to motility arrest by hour 4; however, relative to DMSO-treated control cells, Act. D-treated cell migration speed on 1 kPa was not different until after 8 hours (Δ_8-0_ cell speed on 1 kPa = -4.35 and -3.58 µm/s for DMSO and Act. D, respectively) (Fig. 4B). Transcription inhibition decreased cell spread area by ∼50%, regardless of substrate rigidity (Δ_8-0_ cell area DMSO = +307 and +694 µm^2^ and Δ_8-0_ cell area Act. D = +184 and +318 µm^2^, for 1 and 18 kPa respectively) (Fig. 4C). Circularity exhibited similar trends (Δ_8-0_ cell circularity DMSO = -0.10 and -0.26 and Δ_8-0_ cell circularity Act. D = -0.12 and -0.18, for 1 and 18 kPa respectively) (Fig. 4D).

To investigate the effects of transcriptional feedback blockade on focal adhesion formation and remodeling, we treated cells with Act. D or DMSO at the time of adhesion to 1, 8, or 18 kPa matrices, and stained F-actin with phalloidin and co-immunostained for vinculin at 4 hours post-adhesion (Fig. 4E-G). This represents the time of active matrix mechanotransduction (cf. Figure 2I) and morphodynamic inflection (cf. Figure 4B-D). Acute transcription inhibition increased the total number of vinculin+ adhesion plaques on 1 and 8 kPa matrices (Fig. 4F), but at this time point did not alter the fraction of mature adhesions on any stiffness (Fig. 4G).

**Movies 5, 6: Effects of YAP/TAZ depletion on ECFC migration on soft, moderate, and stiff matrices.** mTomoto-expressing ECFCs were transfected with control (Movie 5) or YAP/TAZ-targeting siRNA (Movie 6). Images were taken in 3-‘minute intervals for 12.5 hours after inhibitor treatment. Time is shown as hours:minutes.

We next tested how matrix mechanical properties altered the dynamics of motile and morphodynamic arrest after YAP/TAZ depletion. To this end, we knocked down expression of both YAP and TAZ, by siRNA transfection at 24 hours prior to trypsinization and adhesion to MeHA hydrogels of 1, 8, or 18 kPa (Supplementary Fig. S2A,B). We then tracked cell migration, morphodynamics, and cytoskeletal and adhesion formation and maturation over 24 hours (Fig. 5A, Supplementary Fig. S2C-H, Movies 5, 6).

**Figure 5:**
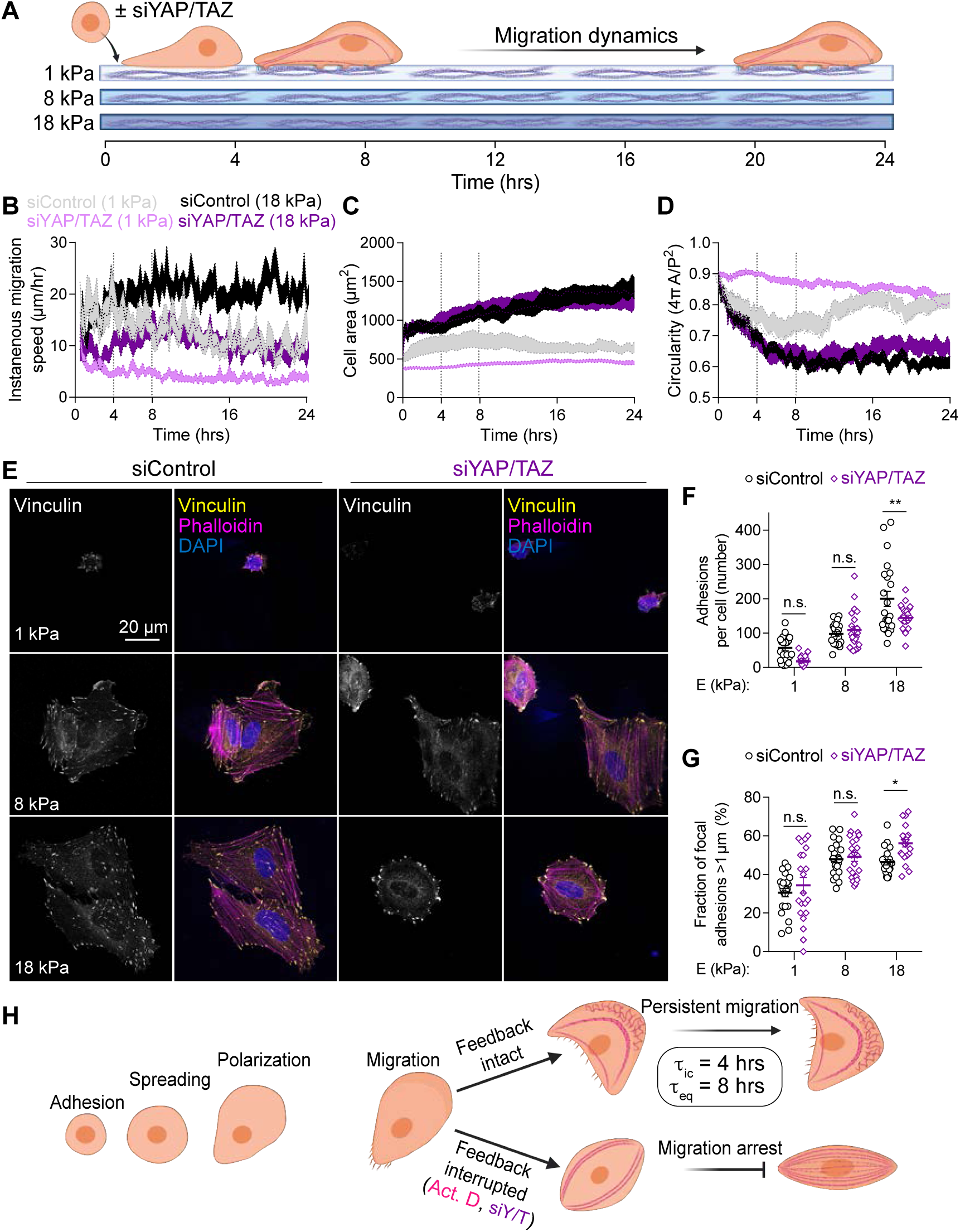
YAP/TAZ-depletion in human ECFCs induces motility and morphodynamic inflection at 4 hours post-attachment, followed by progressive motility arrest, on both soft and stiff substrates. **A.** Experiment schematic: mTomato-expressing ECFCs were treated with siRNA targeting YAP and TAZ or non-targeting control siRNA for 24 hours prior to seeding on 1, 8, or 18 kPa MeHA matrices. Cell morphology and migration were tracked for 24 hours post-attachment or fixed for immunofluorescent imaging at 4 hours post-attachment. **B-D.** Instantaneous migration speed, cell area, and circularity were calculated at 15-minute intervals for 24 hours in four groups: soft-siControl (n = 40 cells), soft-siYAP/TAZ (n = 77 cells), stiff-siControl (n = 48 cells), and stiff-siYAP/TAZ (n = 33 cells). Moderate stiffness groups are shown in Fig. S2. **E.** Immunofluorescence analysis of F-actin (magenta), vinculin (yellow), and nuclei (blue) at 4 hours post-attachment. **F, G.** Quantification of vinculin+ focal adhesions (F) and mature vinculin+ focal adhesions (G), defined as focal adhesions greater than 1 µm. in length. n = 21-22, * p < 0.02, ** p < 0.003, two-way ANOVA with Sidak’s post hoc test. Data are shown as mean ± S.E.M. **H.** Schematic of Adhesion-spreading-polarization-migration assay illustrating persistent migration with intact feedback and migration arrest with feedback interruption by either transcription inhibition (Act. D) or YAP/TAZ depletion (siY/T). Characteristic time scales indicated for initial feedback loop closure (τ_ic_) and motile equilibrium (τ_eq_).

Cell motility and morphodynamics depended on both matrix rigidity and YAP/TAZ feedback. YAP/TAZ depletion reduced cell motility compared to controls on the corresponding substrate, resulting in progressive motility arrest (Fig. 5B). On 1 kPa substrates, YAP/TAZ depletion decreased cell motility continuously with time, reaching a reduced equilibrium migration speed relative to control cells by 8 hours after attachment (Δ_8-0_ cell speed on 1 kPa = -5.37 and -12.96 µm/s for siControl and siYAP/TAZ, respectively). On 18 kPa substrates, YAP/TAZ depletion initially increased migration speed, but transitioned to motility arrest at 8 hours. However, cell motility was lower at every timepoint compared to controls cells on 18 kPa (Δ_8-0_ cell speed on 18 kPa = +12.69 and =+2.97 µm/s for siControl and siYAP/TAZ, respectively). For cells cultured on 1 kPa hydrogels, YAP/TAZ depletion decreased cell spread area by ∼40% at equilibrium (Δ_8-0_ cell area siControl = +213 and +523 µm^2^ and Δ_8-0_ cell area siYAP/TAZ = +132 and +490 µm^2^, for 1 and 18 kPa respectively) (Fig. 5C). Circularity exhibited similar trends (Δ_8-0_ cell circularity siControl = -0.13 and -0.25 and Δ_8-0_ cell circularity siYAP/TAZ = -0.10 and -0.23, for 1 and 18 kPa respectively) (Fig. 5D) Cell area and circularity were not affected by YAP/TAZ inhibition in cells cultured on 18 kPa substrates.

To investigate the role of mechanotransductive feedback in focal adhesion formation and remodeling at the time of active matrix mechanotransduction (cf. Figure 2I) and motility inflection (cf. Figure 5B), we stained F-actin with phalloidin and co-immunostained for vinculin at 4 hours after adhesion to 1, 8, or 18 kPa matrices, with or without YAP/TAZ depletion (Fig. 5E-G). On 18 kPa matrices, YAP/TAZ depletion decreased the total number of vinculin+ adhesion plaques (Fig. 5F), and increased the fraction of adhesions over 1µm in length (Fig. 5G), but did not significantly alter adhesion number or maturity at this time point on either 1 or 8 kPa.

Together, these data indicate that YAP and TAZ transcriptionally regulate the cytoskeleton with a characteristic time scale of ∼4 hours for initial feedback loop closure (τ_ic_), resulting in full motile equilibrium by ∼8 hours (τ_eq_). (Fig. 5H).

To determine whether mechanotransductive feedback kinetics are conserved *in vivo*, we performed two feedback-inhibitor (i.e., Act. D/Puromycin/VP) washout experiments, in which feedback inhibitors were washed out either before (i.e., 3 hrs), or after (i.e., 8 hrs) the characteristic time of initial feedback loop closure (τ_ic_). The 8 hr washout corresponds to our previously-identified time-to-equilibrium (τ_eq_). We hypothesized that, if feedback loop dynamics are conserved, washout at 3 hours would restore morphogenesis while washout after 8 hours would fail to recover.

In the first experiment, we either exposed zebrafish to inhibitors continuously from 29-35 hpf (Fig. 6A), or restored transcriptional feedback at 3 hours by inhibitor washout (Fig. 6B). Without washout, all three drugs significantly slowed ISV growth by 35 hpf; however, washout at 32 hpf significantly restored vessel growth (p = 0.4, 0.12, 0.99 vs. DMSO) (Fig. 6C-G). In contrast, consistent with our hypothesis, washout at 8 hours failed to restore vessel growth kinetics (Fig. 6I-J). These data are consistent with the feedback kinetics observed in human ECFCs *in vitro* and confirm functional lower and upper bounds for YAP/TAZ transcriptional feedback kinetics *in vivo*.

**Figure 6:**
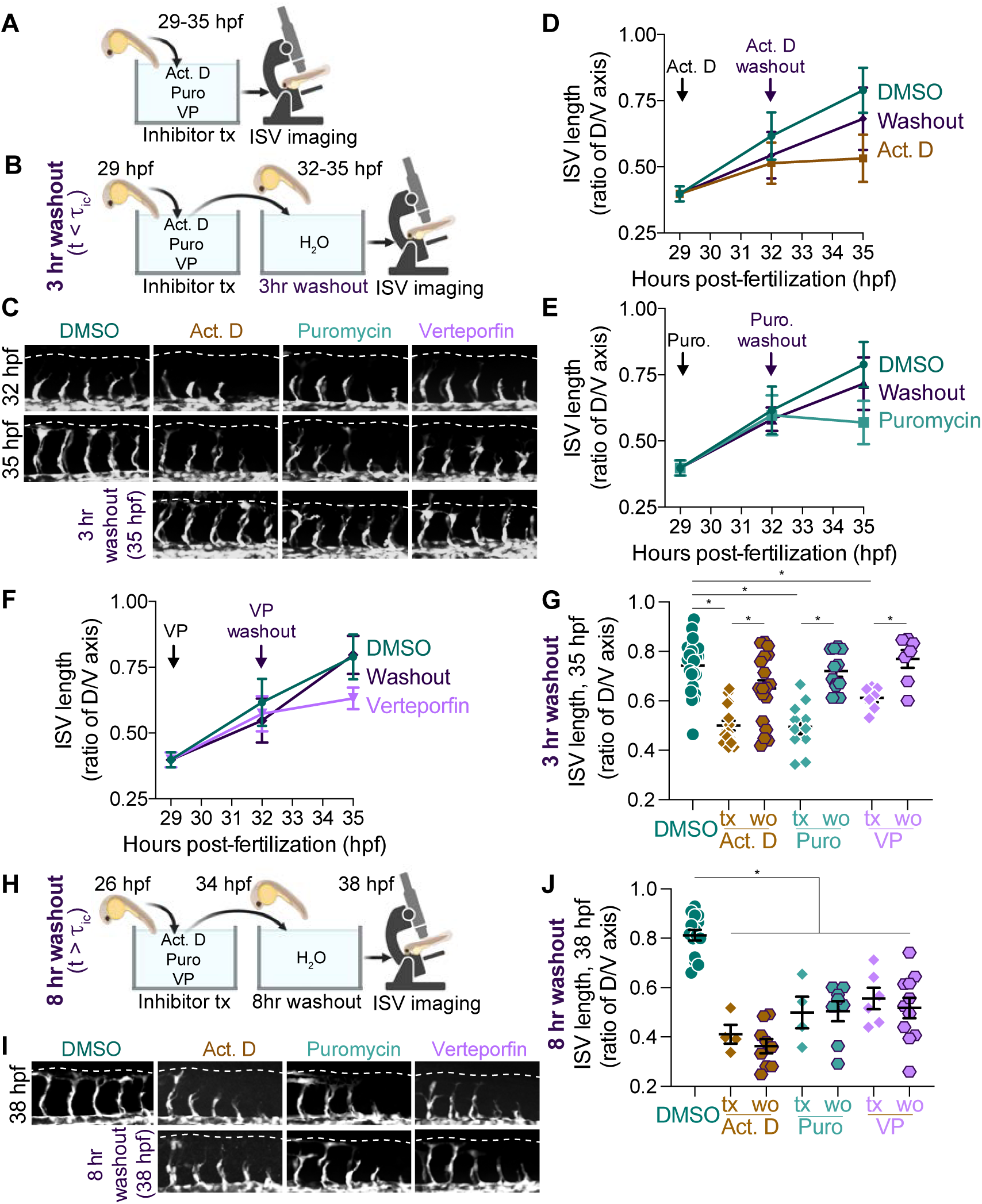
Rescue of vascular morphogenesis by inhibitor washout depends on washout timing. **A.** Schematic diagram of continuous inhibitor treatment. Zebrafish embryos were treated with transcription inhibitor (Act. D), translation inhibitor (Puromycin), or YAP/TAZ inhibitor (Verteporfin), at 29 hpf and imaged at 32 hpf and 35 hpf. **B.** Schematic diagram of three-hour washout, in which inhibitors were washed out after 3 hours, prior to the time to feedback loop closure at τ_ic_ = 4 hrs. Zebrafish embryos were treated with the indicated compounds at 29 hpf, had the inhibitors removed at 32 hpf, and were imaged as the ‘washout’ condition at 35 hpf. **C**. Representative spinning disc confocal images of DMSO, Act. D 25 ug/ml, Puromycin, and Verteporfin treated zebrafish embryos at 32 hpf (top), 35 hpf (middle), or at 35 hpf after 3 hours of incubation in the compounds followed by 3 hours of compound wash out (bottom). **D,E,F**. Quantification of ISV length, tracking individual fish over time. Plots indicate ISV growth rates under conditions of embryos continuously incubated in the indicated compound versus the associated washout condition. Act. D versus washout is shown in (D), Puromycin (Puro) versus washout is shown in (E), and Verteporfin (VP) versus washout shown in (F). **G** Aggregate bulk analysis of zebrafish embryos at isolated time points for each compound versus its associated washout condition. Data are plotted as mean ± S.E.M. DMSO n = 22; Act. D, n = 14 & 18; Puro, n = 11 & 11; and VP, n = 7 & 7 embryos each. * p < 0.05, two-way ANOVA with Sidak’s post hoc test. **H**. Schematic diagrams of our experimental scheme. Zebrafish embryos were treated with the indicated compounds at 26 hpf and imaged at 34 hpf and 38 hpf under conditions of continuous compound incubation versus compound ‘washout’ at 34 hpf. **I**. Representative spinning disc confocal images of DMSO, Act. D 25 ug/ml, Puromycin, and Verteporfin treated zebrafish embryos at 38 hpf either continuously maintained in compound (top) or after 8 hours of incubation in the compounds followed by 4 hours of compound washout (bottom). **J**. Aggregate bulk analysis of zebrafish embryos at 38 hpf for each compound versus its associated washout condition. Data are plotted as mean ± S.E.M. DMSO n = 15; Act. D n = 4 & 9; Puro n = 4 & 8; and VP n = 6 & 11. * p < 0.05, two-way ANOVA with Sidak’s post hoc test. Data points represent the ratio of ISV sprout length at the indicated time point to the total possible sprout length (dotted white line). Five ISVs were averaged per embryo to generate a single data point. A ratio of ‘0’ indicates no ISV formation and ‘1’ indicates completion of ISV morphogenesis.

We next sought to determine how feedback history influences intrinsic state at the time of ECFC attachment to determine cytoskeletal and adhesion dynamics. Unlike acute treatment with Act.D at the time of adhesion (cf. Figure 4), YAP/TAZ depletion by siRNA requires ∼24 hours post-RNAi for protein depletion, but is progressive. Thus, cells trypsinized and replated at 24 hours may already exhibit altered feedback state, analogous to prior studies identifying YAP and TAZ as mediators of cellular mechanical memory (Nasrollahi et al., 2017; Yang et al., 2014a). Therefore, we directly tested the consequences of altering the duration of transcriptional feedback arrest prior to adhesion.

First, we performed either acute or prolonged transcription inhibition (Fig. 7A). Acute transcription inhibition featured Act. D treatment at the time of re-adhesion, as in Figure 4, while prolonged transcription inhibition featured 20 hours of *in situ* Act. D treatment prior to trypsinization and re-adhesion, for a total of 24 hours. We evaluated F-actin and vinculin staining at 4 hours after re-adhesion (Fig. 7B). Prolonged, but not acute, transcription inhibition significantly decreased focal adhesion number at 4 hours after adhesion (Fig. 7C), while both acute and prolonged transcription inhibition significantly increased the fraction of mature focal adhesions (Fig. 7D). Notably, prolonged Act. D treatment produced cells with dense actin stress fibers, robust mature focal adhesions, and decreased spread area after re-adhesion (Fig. 7C-D).

**Figure 7:**
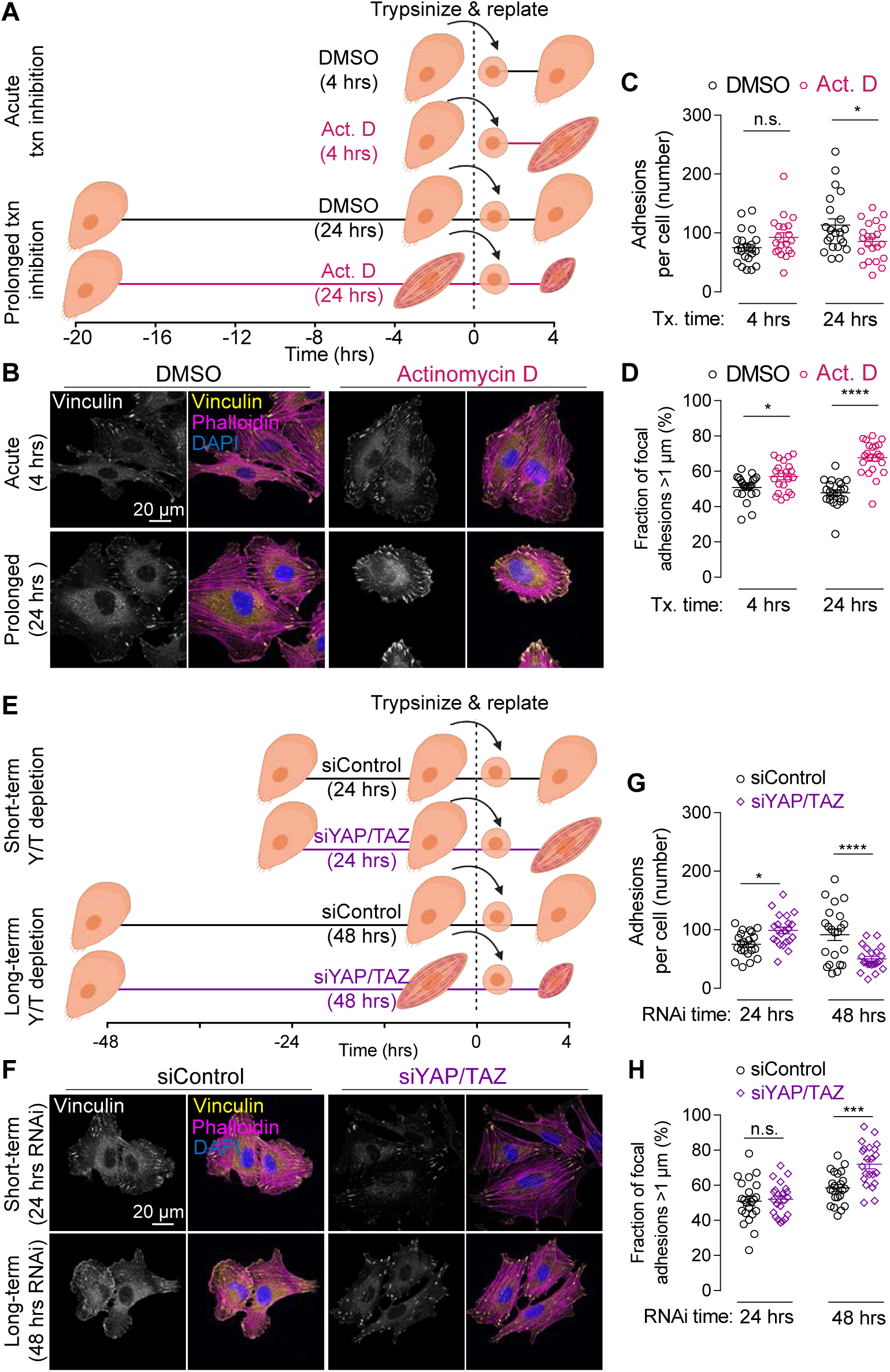
Feedback history and cytoskeletal state prior to re-adhesion alter cytoskeletal and adhesion remodeling. **A.** Transcription inhibition history experiment schematic: mTomato-expressing ECFCs were either pretreated with actinomycin D before plating (−20 hours) or at the time of attachment. All cells were fixed for immunofluorescence at 4 hours after attachment. **B.** Representative immunofluorescent images visualizing actin, vinculin, and nuclei with Alexa Fluor 488-conjugated phalloidin (magenta), Alexa Fluor 594-conjugated secondary (yellow), and DAPI (blue). **C, D.** Vinculin+ focal adhesions (C) and mature vinculin+ focal adhesions (D), defined as greater than 1 µm. n = 20-22, * p < 0.04, **** p < 0.0001, two-way ANOVA with Sidak’s post hoc test. **E.** YAP/TAZ depletion history experiment schematic: mTomato-expressing ECFCs were depleted of YAP and TAZ before plating (24 or 48 hours). All cells were fixed for immunofluorescence at 4 hours after attachment. **F.** Representative immunofluorescent images visualizing actin, vinculin, and nuclei with Alexa Fluor 488-conjugated phalloidin (magenta), Alexa Fluor 594-conjugated secondary (yellow), and DAPI (blue). **G, H.** Vinculin+ focal adhesions (C) and mature vinculin+ focal adhesions (D), defined as greater than 1 µm. n = 22, * p < .03, *** p < 0.0002, **** p < 0.0001 two-way ANOVA with Tukey’s post hoc test.

In parallel, we performed either short-term or long-term YAP/TAZ depletion (Fig. 7E). Short-term YAP/TAZ depletion featured siRNA transfection at 24 hours prior to trypsinization and re-adhesion, as in Figure 5, while long-term transcription inhibition featured siRNA transfection at 48 hours prior to trypsinization and re-adhesion, allowing 24 hours of additional mechanotransductive feedback arrest. We evaluated F-actin and vinculin staining at 4 hours after re-adhesion (Fig. 7F). Short-term YAP/TAZ depletion significantly increased focal adhesion number at 4 hours after adhesion (Fig. 7G). On the other hand, long-term YAP/TAZ depletion significantly decreased focal adhesion number at 4 hours after adhesion (Fig. 7G), but increased the fraction of mature focal adhesions (Fig. 7H). Together, these data suggest that the kinetics of cytoskeletal tension generation and focal adhesion formation and remodeling are dependent on the accumulated cytoskeletal state, modulated by mechanotransductive feedback history.

Next, we observed that *in situ* Actinomycin D treatment for 24 hours increased myosin light chain (MLC) phosphorylation by 3.6-fold (p < 0.04) and promoted myosin association with filamentous structures (Fig. 8A-C), consistent with our prior findings with YAP/TAZ depletion (Mason et al., 2019). To determine how extended transcriptional feedback disruption regulates acute actomyosin activity and function, we performed Actinomycin D treatment for 24 hours or YAP/TAZ depletion for 48 hours, prior to re-adhesion and spreading for 10 minutes. Control cells exhibited minimal stress fibers, with pMLC primarily localized adjacent to the membrane, indicating that trypsinization partially resets the cytoskeleton. In contrast, extended prior transcriptional feedback disruption, by Actinomycin D (Fig. 8D-E) or YAP/TAZ depletion (Fig. 8F-G) prior to trypsinization and adhesion, disrupted cell spreading and exhibited precocious apical myosin activation above the nucleus (Fig. 8D,F y-z projection). To quantify this, we measured the distance of apical pMLC signal above the nucleus (Fig. 8E,G). Prolonged YAP/TAZ depletion had no effect on nuclear height (2.04 vs 2.06 µm; siControl vs. siYAP/TAZ; p > 0.97), but the distance between the top of the nucleus and apical-most pMLC signal increased 3-fold after YAP/TAZ depletion (0.97 vs 3.02 µm; siControl vs. siYAP/TAZ; p < 0.0001). Actinomycin D treatment for 24 hours increased nuclear height 32% (1.73 vs. 2.28 µm; DMSO vs. Act. D; p < 0.007D) and nucleus to apical pMLC distance 59% (1.21 vs. 1.91 µm; DMOS vs. Act. D; p < 0.003). Cross-sectional visualization at the apex of the nucleus in YAP/TAZ inhibited cells revealed pMLC/actin ring accumulation (Fig. 8H).

**Figure 8:**
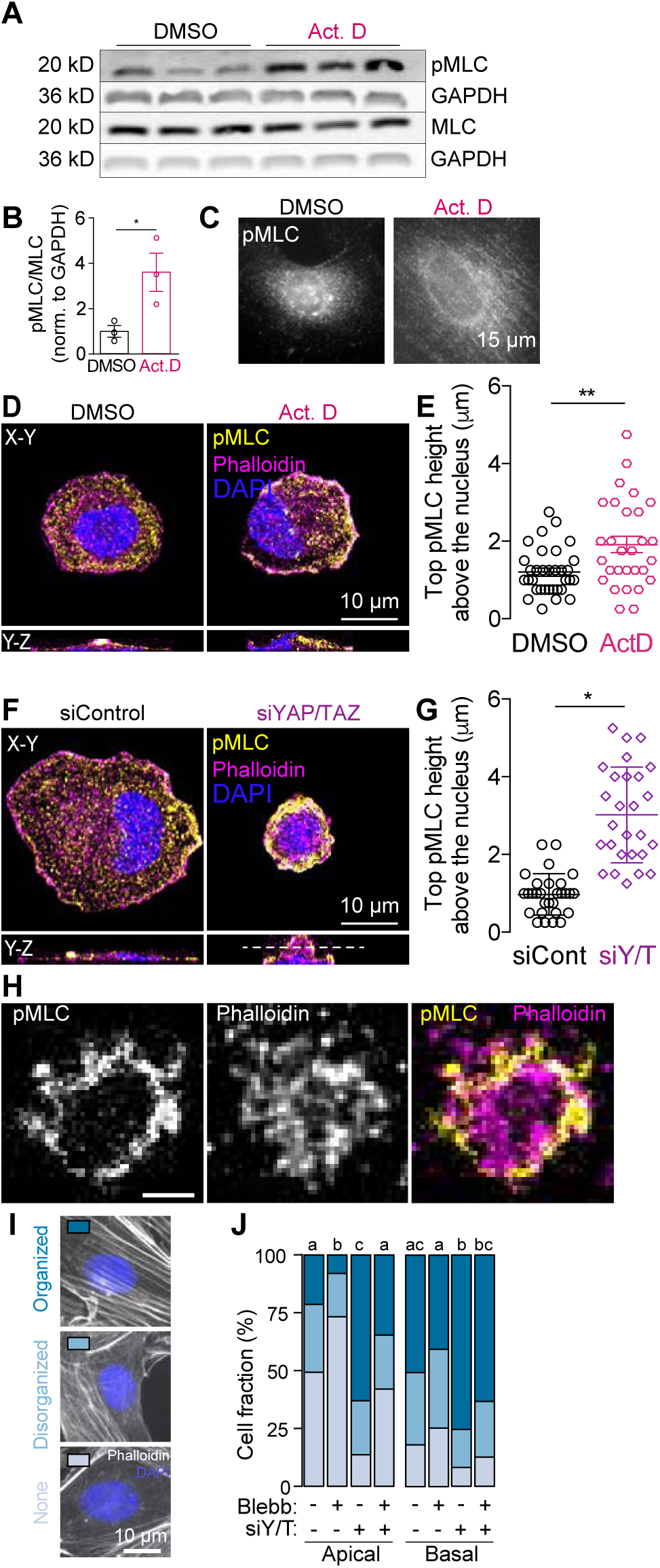
Transcriptional feedback suppresses myosin light chain phosphorylation, limits cell spreading through supranuclear membrane constriction, and promotes actin cap stress fiber maturation. **A-C.** ECFCs were treated for 24 hours with DMSO or Actinomycin D and lysed for relative protein quantification of MLC phosphorylation by immunoblot (A) and immunofluorescent imaging (B, C). Samples used for quantification of pMLC or MLC in (A) were run and transferred in parallel and GAPDH was used as internal loading control so that signal could be normalized between blots. n = 3, * p < 0.05, Student’s two-tailed unpaired t-test. Data shown as mean ± S.E.M. **D-G.** ECFCs were either pretreated with actinomycin D for 24 hours (D, E) or depleted of YAP and TAZ for 48 hours (F, G) prior to plating on glass coverslips. All cells were fixed for 3D confocal imaging at 10 minutes after attachment. **D, F.** Representative immunofluorescent images of basal (x-y) actin, pMLC, and nuclei with Alexa Fluor 647-conjugated phalloidin (magenta), Alexa Fluor 488-conjugated secondary (yellow), and DAPI (blue). Side views are orthogonal projections demonstrating cell height (y-z). **E, G.** Paired measurements of distance of apical-most pMLC signal above the nucleus after Act. D treatment (E) or YAP/TAZ depletion (G). n = 27-33, ** p < 0.003, **** p < 0.0001, student’s two-tailed t-test. Data are shown as mean ± S.E.M. **H.** Representative apical pMLC ring formation adjacent to the cell membrane in Y/T-depleted cell (cross-section corresponds to dotted line in panel H). **I, J.** ECFCs were depleted of YAP and TAZ for 48 hours and treated with low dose (15 µM) Blebbistatin for 1 hour. **I.** Representative immunofluorescent images of actin and nuclei with Alexa Fluor 488-conjugated phalloidin (grey) and DAPI (blue), respectively, depicting ECFC with and without an actin cap. **J.** Number of cells with an organized, disorganized, or no perinuclear actin either on the basal or apical side of the cell. n = 147-150, p < 0.04, Chi square test with Bonferroni’s post hoc test. Data are shown as mean ± S.E.M or as cell number separated into categories.

Myosin activation has been shown to regulate perinuclear actin filament assembly to enable force transmission from focal adhesions to the nucleus (Chambliss et al., 2013), and nuclear strain regulates nuclear permeability, chromatin architecture, and nuclear repositioning during cell motility (Andreu et al., 2021; Elosegui-Artola et al., 2017; Heo et al., 2016; Lee et al., 2007). Therefore, we next tested how YAP/TAZ feedback regulates perinuclear actin filament assembly into ventral stress fibers below the nucleus (basal) and ventral actin cap fibers above the nucleus (apical) (Fig. 8I, J). YAP/TAZ depletion increased the number of cells with an organized apical actin cap 3-fold (p < 0.0001) and basal actin cap by ∼30% (p < 0.0001). To determine whether myosin activity is responsible for the increase in perinuclear actin filaments we treated ECFCs with 15 µM Blebbistatin for 1 hour. This low dose of Blebbistatin has been shown to preferentially target apical actin (Chambliss et al., 2013). Blebbistatin treatment of YAP/TAZ-depleted ECFCs reduced the number of cells with organized apical actin by > 40% (p < 0.0001), without significantly affecting basal actin (p = 0.08). Consistently, YAP/TAZ-mediated transcriptional feedback regulated cell spreading and polarization (Movie 7, 8). Together these data suggest that disrupted transcriptional feedback caused precocious myosin activation to produce apical membrane constriction and alter early cell spreading during peri-nuclear stress fiber polymerization.

**Movie 7: Representative cytoskeletal dynamics of control and Act. D-treated LifeAct-tdTomadto-expressing ECFCs during first hour after adhesion to glass cover slip.** Images were taken in 10-second intervals for 1 hour after attachment. Time is shown as minutes:seconds.

**Movie 8: Representative cytoskeletal dynamics of control and YAP/TAZ depleted LifeAct-tdTomadto-expressing ECFCs during first hour after adhesion to glass cover slip.** Images were taken in 10-second intervals for 1 hour after attachment. Time is shown as minutes:seconds.

### YAP and TAZ regulate transcriptional programs associated with vascular morphogenesis and cytoskeletal regulation

Control and YAP/TAZ depleted cells were seeded on 18kPa MeHa matrices and harvested after four hours to capture early YAP/TAZ-regulated transcriptional changes (Supplementary Table 1). Among the genes most significantly downregulated by YAP/TAZ depletion were *YAP*, TAZ (*WWTR1*), and their known targets (*CTGF*, *CYR61*, *ANKRD1, AXL*) (Dupont et al., 2011; Totaro et al., 2018) (Fig. 9A), suggesting significant functional depletion of YAP and TAZ.

**Figure 9.**
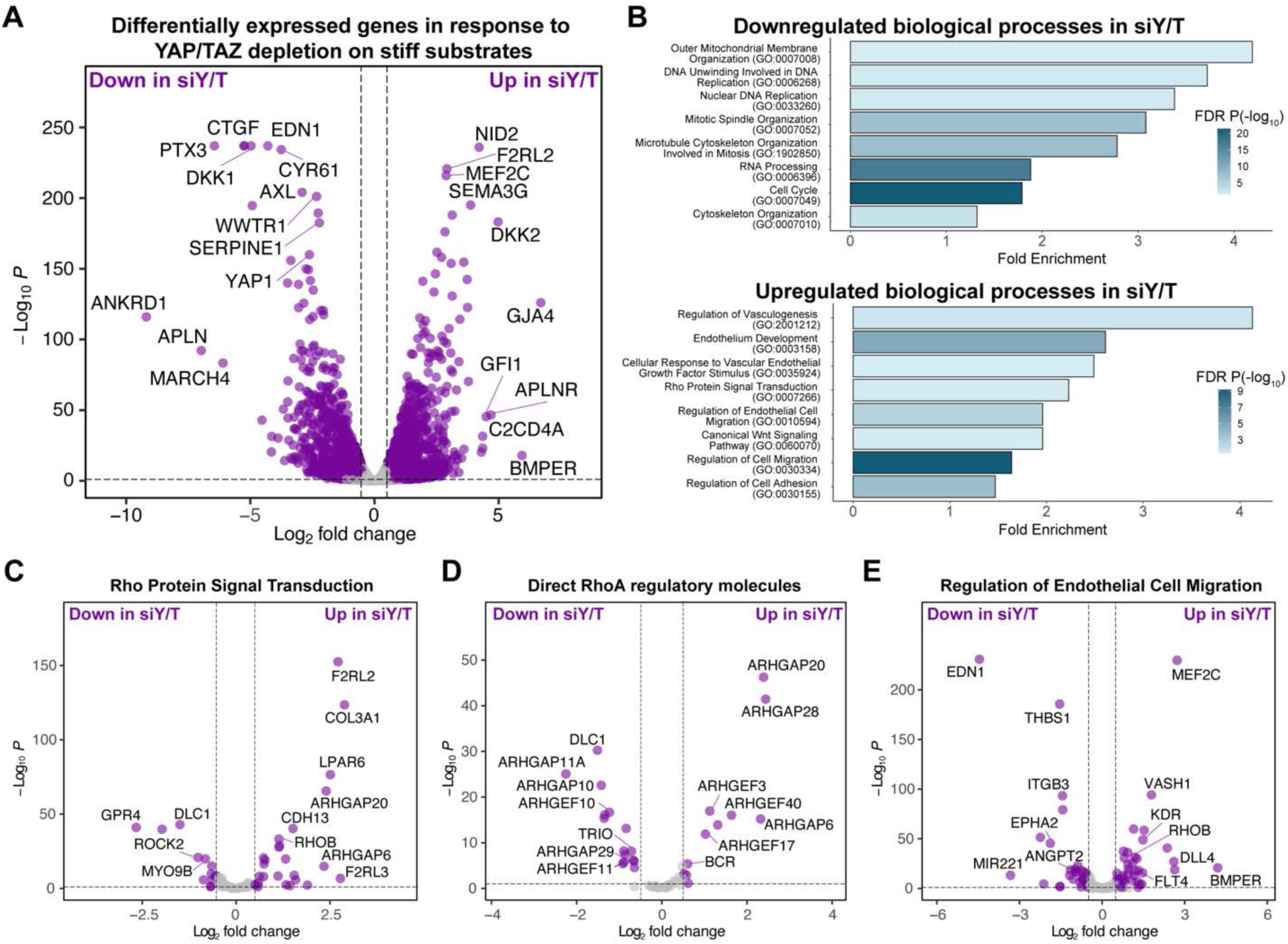
YAP and TAZ regulate transcriptional changes related to vascular development, endothelial cell migration, and RhoA signaling. **A.** Volcano plot of differentially expressed genes (DEGs) from bulk RNA-seq data from cells cultured on 18 kPa substrates. Differential expression analysis was conducted for siYAP/TAZ vs siControl. **B.** Gene Ontology (GO) analysis of significantly enriched biological processes in downregulated (up) or upregulated (down) genes by YAP/TAZ depletion. (PANTHER 17.0 overrepresentation test, p < 0.05, FDR-adjusted Fisher’s exact). **C-D.** Volcano plots of DEGs from siYAP/TAZ vs siControl cells cultured on 18 kPa substrates corresponding to Rho protein signal transduction (GO:0007266) (C), direct RhoA regulatory molecules (compiled from Lawson and Ridley, 2017) (D), and regulation of endothelial cell migration (GO:0010594) (E) (FDR-adjusted Wald test, cutoff criteria p < 0.05 and Log_2_FC > 0.5).

YAP/TAZ depletion caused a similar number of genes to be downregulated (3266 genes) as upregulated (3370 genes) (Fig. 9A). Genes downregulated upon YAP/TAZ depletion were enriched for biological processes including DNA replication, mitosis, RNA processing, and cell cycle (Fig.9B). Genes upregulated upon YAP/TAZ depletion were enriched for biological processes including vasculogenesis, response to VEGF stimulus, Rho protein signal transduction, cell migration, and cell adhesion (Fig. 9B).

Previously, we found that YAP and TAZ mediate negative feedback regulation of the cytoskeleton through transcriptional regulation of RhoA activity. (Mason et al., 2019) We observe this negative regulation in this transcriptomic dataset: YAP/TAZ depletion upregulated genes associated with Rho protein signal transduction (Fig. 9B, C) and altered the expression of genes that directly regulate RhoA activity (Fig. 9D), including previously identified RhoA GAP genes: *DLC1 (ARHGAP7), ARHGAP28*, and *ARHGAP29*. (Mason et al., 2019; van der Stoel et al., 2020)

Although YAP/TAZ depletion/inhibition impaired functional cell motility and vascular morphogenesis (cf. Figures 3, 5), genes upregulated by YAP/TAZ depletion were enriched for GO terms associated with vascular development, cellular response to vascular endothelial growth factor (VEGF) stimulus, and endothelial cell migration (Fig. 9B, E). This appears contradictory but is consistent with our model of YAP/TAZ mechanotransduction as a negative feedback control loop that maintains cytoskeletal homeostasis for persistent endothelial cell motility and blood vessel development. YAP/TAZ depletion upregulated receptors to angiogenic growth factors, including *KDR* and *FLT4,* suggesting YAP/TAZ depleted cells may be more sensitive to VEGF stimulation but remain nonmotile due to cytoskeletal arrest. Together, these data support a role for YAP and TAZ as negative feedback mediators that maintain cytoskeletal homeostasis for endothelial cell migration and vascular morphogenesis.

## Discussion

Here, we show that the transcriptional regulators, YAP and TAZ, execute a conserved mechanotransductive feedback loop that mediates human endothelial cell motility *in vitro* and zebrafish intersegmental vessel morphogenesis *in vivo*. The feedback loop initially closes in 4 hours, achieving cytoskeletal equilibrium by 8 hours, with conserved kinetics *in vivo*. YAP/TAZ transcriptional feedback mediates the cellular response to substrate stiffness mechanosensing and vascular morphogenesis, with mechanotransductive feedback history controlling cytoskeletal and adhesion morphodynamic response to a subsequent mechanical input.

YAP and TAZ are critical regulators of morphogenesis (Dong et al., 2007; Pan, 2010; Phillips et al., 2022), including development of the vertebrate vasculature: YAP and TAZ regulate sprouting angiogenesis and vascular development through both cell-autonomous (Kim et al., 2017; Neto et al., 2017) and cell non-autonomous (Choi et al., 2015; Wang et al., 2017) mechanisms. YAP and TAZ are activated by mechanical cues *in vivo*, including both luminal shear stress and abluminal stretch (Nakajima et al., 2017; Ruehle et al., 2020), and mediate vascular remodeling and low shear stress-induced atherogenesis (Wang et al., 2016). YAP and TAZ also mediate developmental feedback loops. For example, medaka fish embryo morphogenesis against gravitational forces requires YAP/TAZ-mediated transcriptional suppression of RhoA activity through the ARHGAPs (Porazinski et al., 2015).

We identify the mechanotransductive feedback dynamics that mediate human endothelial cell motility *in vitro* and zebrafish vascular morphogenesis *in vivo*. Both broad transcription blockade and targeted YAP/TAZ depletion caused morphodynamic change within 4 hours, altering focal adhesion maturation, followed by motility arrest within 8 hours. Our data suggest that the effectors and kinetics of this feedback loop are conserved in zebrafish. Washout of feedback inhibitors prior to initial feedback loop closure, at 3 hours, restored vessel growth. However, inhibitor washout after 8 hours, longer than the feedback loop timescale, prevented morphogenic rescue. These characteristic time scales are consistent with other studies of human EC motility dynamics in response to diverse mechanical cues (Cai and Schaper, 2008; Weijts et al., 2018). Together, these data suggest a conserved mechanotransductive feedback loop with a characteristic time scale of 4 hours for initial feedback loop closure (τ_ic_), resulting in full motile equilibrium by 8 hours (τ_eq_).

The rate-limiting molecular effectors that determine initial feedback loop completion and equilibrium kinetics are unclear. YAP and TAZ are activated by tension of the actin cytoskeleton and imported into the nucleus from the cytoplasm within minutes through tension-opened nuclear pores (Elosegui-Artola et al., 2017; García-García et al., 2022). Consistent with cytoskeletal remodeling by mechanical perturbation methods (Webster et al., 2014), acute optogenetic activation of either RhoA itself or Rho-activating GEFs induces actomyosin contractility within seconds and YAP nuclear localization within minutes (Berlew et al., 2021; Oakes et al., 2017; Valon et al., 2017). Thus, cytoskeletal activation is likely not rate-limiting. We postulate that the dynamics of YAP/TAZ-dependent transcription and the translation of those target genes are rate-limiting for initial feedback loop completion (τ_ic_ = 4 hours). This is supported by work from us and others in various cell lines showing YAP/TAZ transcriptional responses occur during the first few hours after activation. (Franklin et al., 2020; Mason et al., 2019; Plouffe et al., 2018) The molecular effectors that determine the timescale of new cytoskeletal equilibrium establishment (τ_eq_ = 8 hours) remain unclear.

Our new data support and extend prior models of endothelial cell mechanotransductive feedback by YAP and TAZ (Mason et al., 2019; van der Stoel et al., 2020). However, YAP and TAZ regulation in endothelial cells is context-dependent. Disruption of YAP/TAZ signaling generally inhibits cell motility and angiogenesis (Neto et al., 2018; Ong et al., 2022; Sakabe et al., 2017), but promotes sprouting network formation in the bone marrow endothelium (Sivaraj et al., 2020). We speculate that the role of YAP/TAZ mechanotransductive feedback will be determined by the intrinsic cytoskeletal setpoint in a given endothelial bed. Future research will be necessary to test this hypothesis.

Mechanotransduction is often represented as a one-way path from stimulation to state change, but here we found that mechanotransductive feedback history influences the cytoskeletal response to subsequent stimulation. To directly test the effect of cytoskeletal feedback history, we varied the duration of prior feedback arrest, by either transcription inhibition or YAP/TAZ depletion, and evaluated post-detachment adhesion, spreading, polarization, and migration dynamics. We found that extended feedback arrest prior to reattachment, either by transcription inhibition or YAP/TAZ depletion, altered subsequent cell morphodynamics. This history-dependence is distinct from YAP/TAZ-mediated mechanical memory (Mathur et al., 2020; Nasrollahi et al., 2017; Price et al., 2021; Yang et al., 2014b) but suggests that mechanotransduction is non-linear and features, at minimum, second-order feedback dynamics.

## Limitations

Here we show that both global blockade of de novo gene expression and specific blockade of YAP/TAZ signaling consistently lead to motility arrest *in vitro* and *in vivo*. However, YAP and TAZ are not the only mechanosensitive transcriptional regulators that can modulate the cytoskeleton, and other mechanotransducers are likely to mediate cytoskeletal and morphodynamic feedback. For example, the transcriptional co-activator, MRTF, can also be activated by mechanical cues (Posern et al., 2002), regulates cell migration through transcriptional regulation of cytoskeletal proteins (Leitner et al., 2011), and can co-regulate cytoskeletal feedback in a transcriptional complex with YAP (Katschnig et al., 2017) and/or TAZ (Speight et al., 2016). Future studies will be required to explore roles for this, and other mechanotransducers, in mechanotransductive feedback control of angiogenesis.

Zebrafish embryos enable longitudinal and quantitative imaging of the endothelium during vascular morphogenesis (Phng et al., 2013). Further, embryo transfer between tanks containing inhibitors provides a simple model system for dynamic perturbation of cell signaling. However, every inhibitor has its “off-target” effects, and global inhibition can cause both cell-autonomous and non-cell-autonomous effects on morphogenesis. Additionally, long-term exposure to the pharmacologic inhibitors could impair general cellular function or viability. Future studies will overcome these challenges by orthogonal methods, including inducible and cell-type-specific genetic approaches (Colijn et al., 2022; Pillay et al., 2022) and optogenetic tools (Benman et al., 2022).

## Materials and Methods

### Cell culture and transfection

Human umbilical cord blood-derived ECFCs were isolated at Indiana University School of Medicine and kindly provided by Dr Mervin Yoder. (David A. Ingram et al., 2005; Lin et al., 2023; Rapp et al., 2011; Yoder et al., 2006) ECFCs were cultured as previously described. (David A. Ingram et al., 2005; Mason et al., 2019) Briefly, ECFCs were seeded on collagen (5 μg/cm^2^) coated tissue culture polystyrene (TCPS) and maintained at 37° Celsius and 5% CO_2_ in endothelial growth medium (EGM-2 with bullet kit; Lonza, CC-3162) supplemented with 1% penicillin/streptomycin (Corning) and 10% defined fetal bovine serum (Thermofisher), referred to as full medium. ECFCs were detached from culture dishes using TrypLE Express (Gibco) and used between passages 6 and 8.

pLenti-Lifeact-ubiquitin-tdTomato (Lifeact-tdTomato; addgene: #64048), a filamentous actin-binding peptide labelled with the fluorescent protein tdTomato, was transiently overexpressed in ECFCs under the control of the ubiquitin promoter (Lim et al., 2015). Briefly, ECFCs were plated at a density of 16,000 cells/cm^2^ 24 hours prior to transfection in antibiotic free full media. Lifeact-tdTomato was diluted to 4 ng/µL in EBM2, 105 ng/cm^2^ of DNA per cell culture area. X-tremeGENE™ HP DNA Transfection Reagent (Roche) was used according to the manufacturers protocol, 4µL of transfect reagent was added per 1µg of DNA then incubated for 20-30 minutes at room temperature. Transfection reagent and DNA were then added to ECFCs. ECFCs were used for live imaging 24-72 after transfection.

ECFCs were depleted of YAP and TAZ using custom siRNA (Dharmacon; Table 1) loaded lipofectamine RNAimax (Invitrogen) according to the manufacturer’s instructions, as previously described (Mason et al., 2019). Briefly, ECFCs were seeded on collagen coated 6 well-plates, 10^5^ cells per well, in antibiotic free medium and kept in culture for 24 hours followed by transfection at approximately 50% confluence. Transfection was carried out using a final concentration 0.3% (v/v) lipofectamine RNAimax with 15 pmol RNAi duplexes (custom oligonucleotides; Dharmacon) per well. Transfected ECFCs were used for downstream experiments 24-48 hours post-transfection.

**Table 1:**
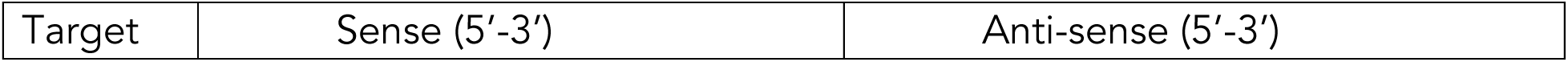
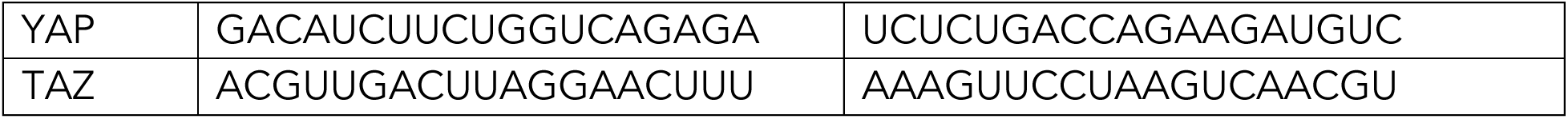
siRNA sense and antisense sequences (Dupont et al Nature 2011, Mason et al JCB 2019)

### MeHA hydrogel synthesis

Methacrylated hyaluronic acid was synthesized as previously described (Vega Ann. Biomed. Eng 2016). Briefly, 1% w/v sodium hyaluronate powder (75 kDa, Lifecore) was reacted with methacrylic anhydride in deionized water (pH 8.5-9) with continuous stirring on ice. MeHA macromer was dialyzed (SpectraPor, 6-8 kDa cutoff) for 5 days then lyophilized for 4 days.

MeHA hydrogels were polymerized on thiolated #1.5 glass coverslips. Coverslips were washed for 20 minutes in 10 M sodium hydroxide (Alfa Aesar) then washed twice in deionized water and dried. Dried coverslips were incubated in toluene (Fisher Scientific) with 12.5% v/v (3-mercaptopropyl) trimethoxysilane (Sigma Aldrich) and 4.2% v/v hexylamine (Sigma Aldrich) for 1 hour. Thiolated coverslips were washed twice in toluene, dried at 100° C, and stored under nitrogen for up to one month prior to use.

MeHA hydrogels were formed from a 3% w/v MeHA macromer solution functionalized with 1 mM RGD peptide (GenScript) in 0.2 M Triethanolamine (TEOA; Sigma Aldrich) buffer for 30 minutes at room temperature. The dithiol cross-linker Dithiothreitol (DTT; Thermo Scientific) was added to the macromer solution at concentrations of 2.4, 4.8, or 9.6 mM. Sufficient volume to produce 60 µm thick hydrogels were pipetted onto thiolated coverslips and flattened with Rain X treated hydrophobic coverslips. MeHA was polymerized at 37° C for 3 hours then washed PBS and stored at 4° C for cell culture.

MeHA hydrogel elastic modulus was measured using atomic force microscopy (AFM; Bruker Bioscope Catalyst). A 1 µm diameter SiO_2_ spherical probe (∼0.068 N/m stiff; Novascan) was indented ∼0.5 µm to generate a force-displacement curve. Average elastic modulus was calculated using the hertz contact model formula. Measurements were taken from 10 points for each sample, four samples were measured per hydrogel formulation.

### Immunofluorescence

Cells were washed twice in EBM-2 and fixed in 4% paraformaldehyde (Alfa Aesar) diluted in EBM-2 for 15 minutes at room temperature. Cells were permeabilized for 5 minutes with 0.1% triton x-100 (amresco) and blocked in PBS containing with 5% goat serum (Cell Signaling) for one hour. Fixed samples were incubated with primary antibodies diluted in PBS with 1% goat serum and .1% tween-20: YAP (1:200, Cell Signaling, 14074), TAZ (1:250, Cell Signaling, 4883), Vinculin (1:100, Sigma, V9131), pMLC (1:400, abcam, ab2480), pMLC (1:200, Cell signaling, 3675). Primary antibodies were detected secondaries antibodies diluted in PBS with 1% goat serum: polyclonal Alexafluor 594-conjugated anti-rabbit IgG (1:1000, Cell Signaling, 8889), and polyclonal Alexafluor 488-conjugated anti-mouse IgG (1:1000, Cell Signaling, 4408). F-actin was stained using Alexa fluor 488, 594, or 647-conjugated phalloidin (1-3 unit/mL; Life Technologies) 30 minutes. Nuclei were stained with DAPI (Sigma Aldrich) diluted 1:1000 for 30 minutes. Samples were mounted in ProLong® Gold Antifade solution (Thermo Fisher Scientific).

### ECFC live migration

ECFC motility was tracked live on MeHA by fitting custom circular PDMS (Dow Corning; 1.75 cm diameter) wells to thiolated coverslips using PDMS to bond the coverslips and glass. Hydrogels were formed in the wells as described above and used for live imaging. For imaging cell attachment mTomato-expressing ECFCs were plated 1-2,000 cells/cm^2^ on MeHA and imaged live using a Zeiss Axio Observer inverted microscope with an automated stage for 4 hours at 3-minute intervals followed by 20 hours at 15 minute intervals. Cells were tracked across 5 ROI’s for each substrate 5-15 cells per ROI.

For continuously attached cells treated with inhibitors mTomato-expressing ECFCs were first attached for 24-48 hours to collagen coated glass chamber slides before imaging for 24-36 hours at 15-minute intervals. Actinomycin D (Sigma Aldrich), Puromycin (Takara Bio) or DMSO (Sigma Aldrich) were added at the indicated times. During live imaging cells were maintained at 37° C, 5% CO^2^, 95% relative humidity using a stage top type incubation chamber (Heating insert P; Pecon). For all live imaging experiments migration was tracked in 5 regions per condition. For imaging experiments longer than 1 hour a layer of silicone oil (ibidi) was added on top of culture wells to limit evaporation. Cells were tracked across 5 ROI’s 25-50 cells per ROI.

Morphological and positional tracking of individual cells was performed using the ADAPT plugin for the open source image analysis software FIJI (Barry et al., 2015). Using ADAPT, the cell outlines were thresholded either using the Li, Huang, or Default thresholding method, depending on tracking fidelity. Objects below a realistic size threshold (200 or 400 µm^2^, depending on the experiment) or that could not be tracked for > 80% of the imaging time were not recorded. Object smoothing was set to .5 and 4 erosion iterations were used to accurately track cell boundaries as a function of time.

### Lamellipodia tracking

ECFCs expressing Lifeact-tdTomato were trypsinized and maintained in suspension in full media in the presence of either DMSO, Actinomycin D, or puromycin (Takara Bio) for 5 minutes then reattached to collagen-coated coverslips. Alternatively, ECFCs treated with siRNA for 24 hours were then transfected with Lifeact-tdTomato for 24 hours then reattached to collagen-coated coverslips. Lifeact-expressing ECFCs were imaged using a 63x objective (NA:1.2) in 10s intervals for 1 hour on a Zeiss axio observer. Lifeact signal-to-noise ratio was improved by two sequential rounds of thresholding and variance filtering (in 10 and 4 pixel interrogation windows). Videos were processed in matlab to track lamellipodial dynamics 5 µm from the edge of the cell. Background was subtracted and lamellipodia displacements (d) tracked using the built-in Farneback optic flow function in matlab. The cell centroid at each time was used as an internal reference point for constructing a position matrix (p). The dot product (d·p) of the position matrix (p) and the displacement matrix was then used to classify lamellipodia as protrusive (d·p > 0) or retractive (d·p < 0). Average lamellipodia protrusion or retraction velocity were calculated across 20 minutes of cell spreading. Experiments were repeated in triplicate where at least 1-4 cells were images per condition.

### Microscopy and image analysis

Epifluorescence images of fixed and live samples were taken on Zeiss Axio Observer equipped with a monochromatic Axiocam 702 (Zeiss) at 23° C using 5x (NA: 0.16), 10x (NA: 0.3), 20x (NA: 0.8), 40x (NA: 0.6), and 63x (NA: 1.2) Zeiss objectives. Data acquisition was done using the ZEN imaging suite (Zeiss). Confocal image stacks were taken on a laser-scanning Leica DMI 6000 inverted microscope using a 63x (NA: 1.4) Leica objective. Deconvolution of z-stacks were performed using Hugyens professional image processing application with a theoretical point spread function.

Focal adhesion morphology was measured after rolling ball subtraction with a radius of 50 pixels and thresholding the Huang thresholding method. Adhesions were identified on a per cell basis, across three experiments as being greater than .1 µm^2^ and less than 10 µm^2^. All image analysis (morphometrics, image adjustments, and individual cell tracking) were performed using an open access NIH software platform, FIJI (Schindelin Nat Methods 2012).

Z-stacks of pMLC, actin, and DAPI signal were taken through the thickness of a cell at a depth of 0.75 µm. Images underwent deconvolution using Hugyens professional described above and height was measured using the z-axis intensity profile tool from FIJI. DAPI and pMLC height were estimated as the position above the nuclei with less than 10 % of the max fluorescent intensity of a given stack, where the region with the highest fluorescent intensity was typically in line with the middle of the nucleus. Representative line plots show relative fluorescent intensity, normalized across a given image stack. Fluorescent signal less than 10 % of the max fluorescent intensity was assumed to be background and made zero.

### Quantitative reverse transcription PCR (RT-qPCR)

Total RNA was isolated and purified using the RNeasy mini kit (Qiagen). 0.5 µg of total RNA was reversed transcribed (Applied Biosystems) using the manufacturer’s instructions in a Mastercycler nexus gradient (Eppendorf). cDNA was mixed with SYBR green mastermix (Applied Biosystems) and 0.4 µM forward and reverse primers (CTGF: TTAAGAAGGGCAAAAAGTGC (Forward 5’-3’) and CATACTCCACAGAATTTAGCTC (Reverse 5’-3’); Sigma Aldrich) in wells of 96 well PCR plate (Applied Biosystems). cDNA was amplified and quantified using a Step-one Plus real-time PCR system (Applied Biosystems). Relative gene expression was quantified using the ΔΔC_T_ method normalizing target gene expression with the housekeeping gene 18S.

### RNA-sequencing & Gene Ontology analysis

Total RNA was assessed for quantity and quality with RNA ScreenTape assay of Agilent 2200 TapeStation System (Agilent Technologies). Average RIN values were 10. 200 ng of RNA was used as input for library prep. Libraries were prepared using TruSeq Stranded mRNA HT Sample Prep Kit (Illumina) as per standard protocol in the kit’s sample prep guide. Libraries were assayed for overall quality using D1000 ScreenTape assay of Agilent 2200 TapeStation System (Agilent Technologies). Average library molarity from Tapestation was 103.331+/-24.6 nM. Samples were multiplexed for sequencing. A MiSeq test lane was run to check pool balance followed by 100bp single-read sequencing on an Illumina HiSeq 4000 sequencer. Illumina’s bcl2fastq2 version v2.20.0.422 software was used for demultiplexing and converting bcl to fastq files.

Reads were filtered to retain only high-quality reads. In addition, ribosomal reads and repeats were eliminated by alignment to a generic set of ribosomal/repeat sequences. Remaining reads were processed with RNA-Seq Unified Mapper (RUM) (Grant et al., 2011). Following RUM alignments to human genome (build hg19) and known transcripts (RefSeq, UCSC, ENSEMBL), a feature-level quantitation (transcript, exon, and intron) were output as result. To analyze global gene expression profiles, the number of uniquely aligning reads to mRNA transcripts in RefSeq were extracted from the RUM output.

Statistically significant differences and fold changes in individual genes were computed via DESEq2 v. 1.36.0. DEGs were defined as having a Wald test FDR-adjusted P value less than 0.05. GO was conducted separately for genes significantly upregulated or downregulated in siYAP/TAZ samples compared to siControl using PANTHER overrepresentation test on PANTHER 17.0, available on the GO Resource. We used the GO biological process complete annotation set with Fisher’s exact test and FDR-corrected P values. The test was applied using a query set of genes significantly elevated or decreased (Wald test FDR-adjusted P value less than 0.05; n = 3370 upregulated genes and n = 3266 downregulated genes in siYAP/TAZ) and a background of all nonzero features identified in the bulk transcriptome.

The raw RNA sequencing data, DESEq2 results, and enrichment analysis are available in Supplementary Table 1.

### Fluorescent western blot

Briefly, cells were lysed in RIPA buffer (Cell Signaling) and mixed with LDS loading buffer (Invitrogen) and sample reducing agent (Invitrogen) reduced at 70°C for 10 minutes. Samples were processed for gel electrophoresis on 4-12% NuPAGE Bis-Tris gels (Invitrogen) in either MES or MOPS NuPAGE running buffer (Invitrogen). Proteins were blotted on low-fluorescent PVDF (Biorad) with NuPAGE transfer buffer (Invitrogen), 20% methanol (Fisher Scientific), 0.1% v/v antioxidant (Invitrogen). PVDF was blocked with TBS blocking buffer (LI-COR biosciences). Primary antibodies: YAP (1:500, Cell Signaling, 14074), TAZ (1:1000, Cell Signaling, 4883), pMLC (1:1000, abcam, ab2480), MLC (1:200; Santa Cruz; sc-28329), GAPDH (1:3000, Cell Signaling, 5174) were incubated overnight in blocking buffer with .2% tween-20 (Fisher Scientific). Primary antibodies were identified by IRDye 800 or 680-conjugated secondaries (1:20,000; LI-COR biosciences; 926-32212 & 926-68073) incubated in blocking buffer with 0.2% tween-20 and 0.02% SDS (Amresco). Fluorescent western blots were imaged using a LI-COR odyssey imager (LI-COR biosciences). Quantification was performed using Image J were target protein expression was normalized to GAPDH expression. For relative MLC phosphorylation quantification samples were run and transferred in parallel. MLC or pMLC signal was first normalized to GADPH signal then pMLC was normalized to total MLC.

### Zebrafish husbandry, treatments, and imaging

Zebrafish *(Danio rerio)* embryos were raised and maintained as described (Kimmel et al., 1995). Zebrafish husbandry and research protocols were reviewed and approved by the Washington University in St. Louis Animal Care and Use Committee. The zebrafish transgenic line Tg*(fli:egfp) ^y1^* is previously published (Lawson et al., 2002).

Pharmacologic treatments of zebrafish embryos were done by adding all small molecule inhibitors to zebrafish embryos housed off system in petri dishes. Embryos were allowed to undergo gastrulation and the inhibitors were added beginning at 28-29 hours post fertilization. The following inhibitors were resuspended in DMSO and used at the final concentration as indicated: Actinomycin D (1.0 and 2.5 µg/ml; Tocris; 1229); Puromycin (10ug/ml; Tocris; 4089); Y27632 (10uM; Tocris; 1254); Rockout (50uM; Sigma Aldrich; 555553); Blebbistatin (10uM; Tocris; 1760); Verteporfin (10uM; Sigma Aldrich; SML0534); BDM (2,3-Butanedione monoxime; 20mM; Sigma Aldrich; B0753).

Fluorescent images were collected utilizing a Nikon Ti2 and Yokogawa CSU-W1 spinning disk confocal microscope between 28-48 hpf at 10x magnification. To maintain timing consistency between treatments, embryos were fixed in 4% PFA for 24 hours at 4°C before being embedded in 0.8% low melting point agarose for imaging. Z-stacks were acquired using a 1um step size, and max projections generated using FIJI software. ISV length was measured using FIJI and defined as the length of the ISV bounded between the dorsal aorta and the DLAV (dorsal longitudinal anastomotic vessel). Five ISV’s were measured per embryo and an average per embryo is represented in the graphs. The data is normalized to the height of the dorsal ventral axis of the animal it was collected from to account for differences is mounting and imaging.

## Statistics

Statistical analyses of cell migration and ISV morphogenesis were performed on Graphpad Prism 6 statistical analysis package. Data are presented with data points were possible and mean ± standard error or standard deviation; figure captions describe data presentation. All experiments were performed at least in triplicate. Multiple comparisons were made using analysis of variance (ANOVA) with Tukey or Sidak’s *post hoc* test for pairwise comparisons of normally distributed homoscedastic data. Data were considered to fit the ANOVA assumptions if the residuals of a data set were normally distributed. Data that did not meet the ANOVA criteria were analyzed by Kruskal-Wallis with Dunn’s *post hoc* test. Comparisons between two data-sets were made using Student’s unpaired two-tailed t-test, or non-parametric by Mann-Whitney, when necessary.

## Supporting information

RNAseq Data

## Acknowledgements

This work was supported by the National Institutes of Health: R01 AR073809 (to JDB), R01 GM143400 (to JDB, BYC), and the National Science Foundation Science and Technology Center CMMI 1548571 (to JDB). Figures include images constructed, in part, using Biorender. We gratefully acknowledge insightful questions and suggestions from Lance Davidson, Univ. of Pittsburgh.

## Supplementary Data

**Supplementary Figure 1.**
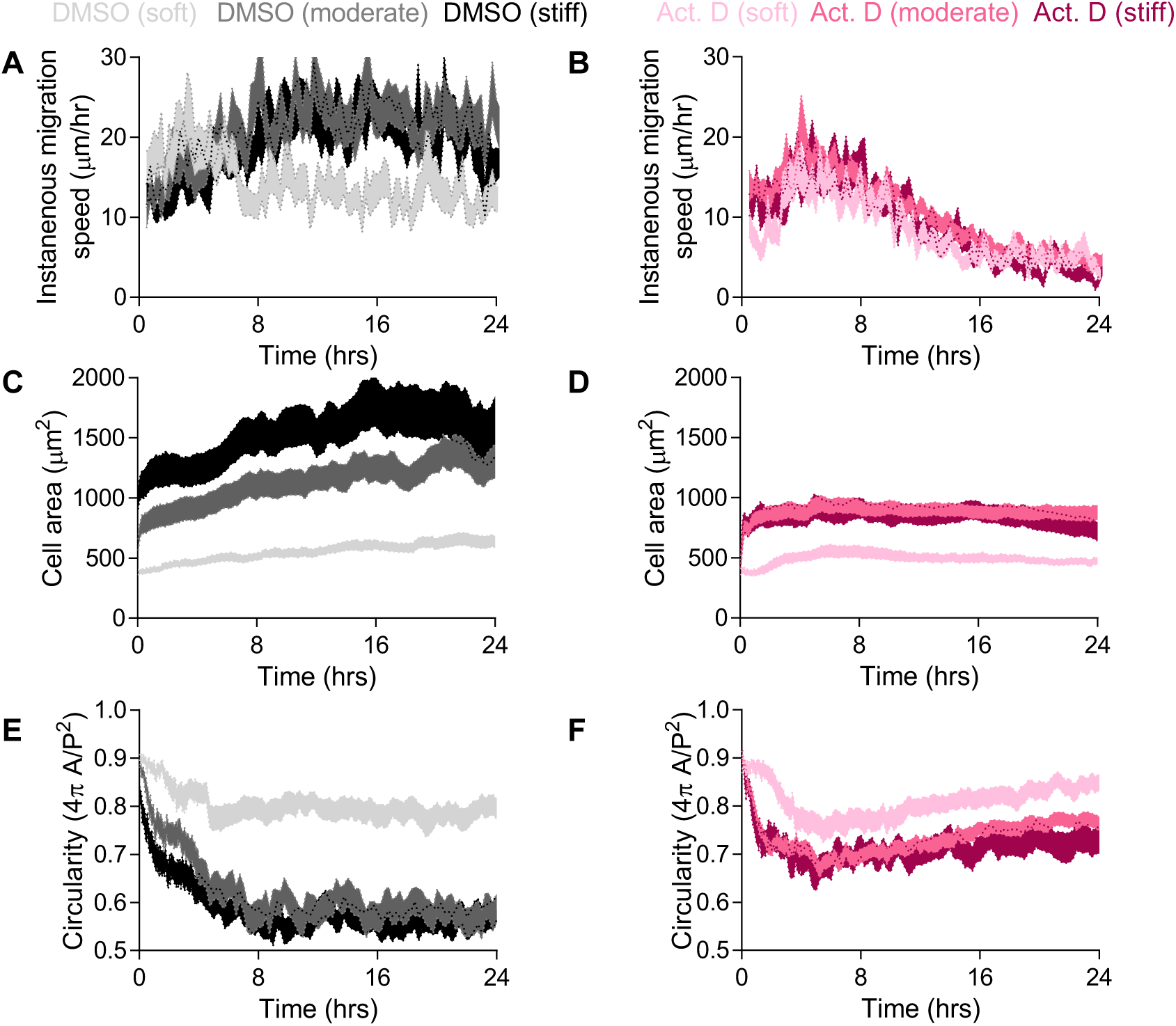
Effects of transcription inhibition on ECFCs motility and morphodynamics after attachment to soft (1 kPa), moderate (8 kPa), and stiff (18 kPa) matrices. (A-E) mTomato-expressing ECFCs were seeded on hydrogels with either DMSO or actinomycin D (0.1 µg/mL) and cell (A, B) motility, (C, D) area, and (E, F) circularity tracked as a function of time after attachment. Soft-DMSO (n = 48 cells), soft-Act.D (n = 63 cells), Moderate-DMSO (n = 97 cells), Moderate-Act.D (n = 39 cells), stiff-DMSO (n = 41 cells), and stiff-Act.D (n = 36 cells). Figures include soft and stiff data reproduced from Figure 4, for comparison. Data are shown as mean ± S.E.M.

**Supplementary Figure 2:**
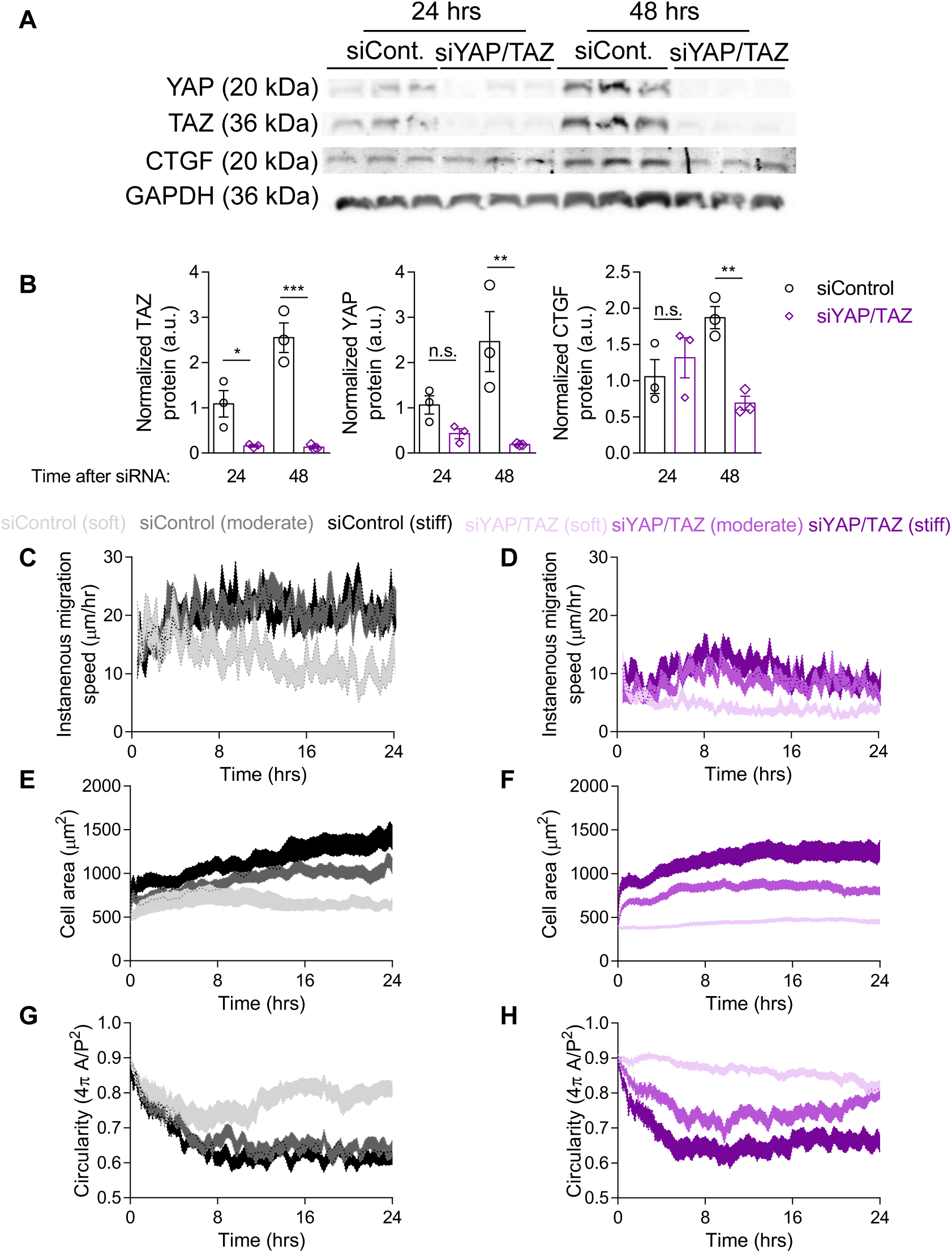
Effects of YAP/TAZ depletion on ECFCs motility and morphodynamics after attachment to soft (1 kPa), moderate (8 kPa), and stiff (18 kPa) matrices. (A-B) ECFCs were depleted of YAP and TAZ for 24 or 48 hours then lysed for relative protein quantification by Western Blot. n = 3, * p < 0.04, ** p < 0.007, *** p = 0.0001, one-way ANOVA with Sidak’s post-hoc test. Data are shown as mean ± S.E.M. (C-H) mTomato-expressing ECFCs depleted of YAP and TAZ were seeded on hydrogels and cell (C, D) motility, (E, F) area, and (G, H) circularity tracked as a function of time after attachment. Soft-siControl (n = 40 cells), soft-YAP/TAZ (n = 77 cells), Moderate-siControl (n = 47 cells), Moderate-siYAP/TAZ (n = 57 cells), stiff-Control (n = 48 cells), and stiff-siYAP/TAZ (n = 33 cells). Figures include soft and stiff data reproduced from Figure 5, for comparison. Data are shown as mean ± S.E.M.

**Supplementary Table 1. Bulk RNA sequencing analysis from ECFCs cultured on 18 kPa substrates. 1a.** Raw data in reads from control (C_st, n=2) or YAP/TAZ depleted (YT_st, n=4) ECFCs cultured in 18 kPA substrates. **1b.** Differential gene expression analysis for siYAP/TAZ vs siControl samples computed via DESEq2. **1c.** Enrichment analysis of biological processes from downregulated genes by YAP/TAZ depletion. **1d.** Enrichment analysis of biological processes from upregulated genes by YAP/TAZ depletion.

## Notes

### Competing Interest Statement

The authors have declared no competing interest.

### Summary of Updates

Manuscript revised after Peer Review at eLife.

## References Cited

Andreu I, Granero-Moya I, Chahare NR, Clein K, Jordàn MM, Beedle AEM, Elosegui-Artola A, Trepat X, Raveh B, Roca-Cusachs P. 2021. Mechanosensitivity of nucleocytoplasmic transport. bioRxiv 2021.07.23.453478. doi:10.1101/2021.07.23.453478

Barry DJ, Durkin CH, Abella J V, Way M. 2015. Open source software for quantification of cell migration, protrusions, and fluorescence intensities. The Journal of cell biology 209:163–80. doi:10.1083/jcb.201501081

Benecke B-J, Ben-Ze’ev A, Penman S. 1978. The control of mRNA production, translation and turnover in suspended and reattached anchorage-dependent fibroblasts. Cell 14:931– 939. doi:10.1016/0092-8674(78)90347-1

Benman W, Berlew EE, Deng H, Parker C, Kuznetsov IA, Lim B, Siekmann AF, Chow BY, Bugaj LJ. 2022. Temperature-responsive optogenetic probes of cell signaling. Nature Chemical Biology 18:152–160. doi:10.1038/s41589-021-00917-0

Berlew EE, Kuznetsov IA, Yamada K, Bugaj LJ, Boerckel JD, Chow BY. 2021. Single-Component Optogenetic Tools for Inducible RhoA GTPase Signaling. Advanced Biology 5. doi:10.1002/adbi.202100810

Boselli F, Freund JB, Vermot J. 2015. Blood flow mechanics in cardiovascular development. Cellular and Molecular Life Sciences 72:2545–2559. doi:10.1007/s00018-015-1885-3

Cai W, Schaper W. 2008. Mechanisms of arteriogenesis. Acta biochimica et biophysica Sinica 40:681–92.

Chambliss AB, Khatau SB, Erdenberger N, Robinson DK, Hodzic D, Longmore GD, Wirtz D. 2013. The LINC-anchored actin cap connects the extracellular milieu to the nucleus for ultrafast mechanotransduction. Scientific reports 3:1087. doi:10.1038/srep01087

Choi H-J, Zhang H, Park H, Choi K-S, Lee H-W, Agrawal V, Kim Y-M, Kwon Y-G. 2015. Yes-associated protein regulates endothelial cell contact-mediated expression of angiopoietin-2. Nature communications 6:6943. doi:10.1038/ncomms7943

Colijn S, Yin Y, Stratman AN. 2022. High-throughput methodology to identify CRISPR-generated Danio rerio mutants using fragment analysis with unmodified PCR products. Developmental Biology 484:22–29. doi:10.1016/J.YDBIO.2022.02.003

Cosgrove BD, Mui KL, Driscoll TP, Caliari SR, Mehta KD, Assoian RK, Burdick JA, Mauck RL. 2016. N-cadherin adhesive interactions modulate matrix mechanosensing and fate commitment of mesenchymal stem cells. Nature materials 15:1297–1306. doi:10.1038/nmat4725

Dong J, Feldmann G, Huang J, Wu S, Zhang N, Comerford SA, Gayyed MF, Anders RA, Maitra A, Pan D. 2007. Elucidation of a universal size-control mechanism in Drosophila and mammals. Cell 130:1120–33. doi:10.1016/j.cell.2007.07.019

Dupont S, Morsut L, Aragona M, Enzo E, Giulitti S, Cordenonsi M, Zanconato F, Le Digabel J, Forcato M, Bicciato S, Elvassore N, Piccolo S. 2011. Role of YAP/TAZ in mechanotransduction. Nature 474:179–183. doi:10.1038/nature10137

Elosegui-Artola A, Andreu I, Beedle AEM, Lezamiz A, Uroz M, Kosmalska AJ, Oria R, Kechagia JZ, Rico-Lastres P, Le Roux A-L, Shanahan CM, Trepat X, Navajas D, Garcia-Manyes S, Roca-Cusachs P. 2017. Force Triggers YAP Nuclear Entry by Regulating Transport across Nuclear Pores. Cell 171:1397–1410.e14. doi:10.1016/J.CELL.2017.10.008

García-García M, Sánchez-Perales S, Jarabo P, Calvo E, Huyton T, Fu L, Ng SC, Sotodosos-Alonso L, Vázquez J, Casas-Tintó S, Görlich D, Echarri A, Del Pozo MA. 2022. Mechanical control of nuclear import by Importin-7 is regulated by its dominant cargo YAP. Nature Communications 13:1174. doi:10.1038/s41467-022-28693-y

Heo S-J, Driscoll TP, Thorpe SD, Nerurkar NL, Baker BM, Yang MT, Chen CS, Lee DA, Mauck RL. 2016. Differentiation alters stem cell nuclear architecture, mechanics, and mechano-sensitivity. eLife 5. doi:10.7554/eLife.18207

Huynh J, Nishimura N, Rana K, Peloquin JM, Califano JP, Montague CR, King MR, Schaffer CB, Reinhart-King CA. 2011. Age-related intimal stiffening enhances endothelial permeability and leukocyte transmigration. Science translational medicine 3:112ra122. doi:10.1126/scitranslmed.3002761

Ingram David A., Caplice NM, Yoder MC. 2005. Unresolved questions, changing definitions, and novel paradigms for defining endothelial progenitor cells. Blood 106:1525–1531. doi:10.1182/blood-2005-04-1509

Ingram David A, Mead LE, Moore DB, Woodard W, Fenoglio A, Yoder MC. 2005. Vessel wall-derived endothelial cells rapidly proliferate because they contain a complete hierarchy of endothelial progenitor cells. Blood 105:2783–6. doi:10.1182/blood-2004-08-3057

Ingram DA, Mead LE, Tanaka H, Meade V, Fenoglio A, Mortell K, Pollok K, Ferkowicz MJ, Gilley D, Yoder MC. 2004. Identification of a novel hierarchy of endothelial progenitor cells using human peripheral and umbilical cord blood. Blood 104:2752–60. doi:10.1182/blood-2004-04-1396

Katschnig AM, Kauer MO, Schwentner R, Tomazou EM, Mutz CN, Linder M, Sibilia M, Alonso J, Aryee DNT, Kovar H. 2017. EWS-FLI1 perturbs MRTFB/YAP-1/TEAD target gene regulation inhibiting cytoskeletal autoregulatory feedback in Ewing sarcoma. Oncogene 36:5995–6005. doi:10.1038/onc.2017.202

Kim Jongshin, Kim YH, Kim Jaeryung, Park DY, Bae H, Lee D-H, Kim KH, Hong SP, Jang SP, Kubota Y, Kwon Y-G, Lim D-S, Koh GY. 2017. YAP/TAZ regulates sprouting angiogenesis and vascular barrier maturation. The Journal of Clinical Investigation 127:3441–3461. doi:10.1172/JCI93825

Kimmel CB, Ballard WW, Kimmel SR, Ullmann B, Schilling TF. 1995. Stages of embryonic development of the zebrafish. Developmental Dynamics 203:253–310. doi:10.1002/aja.1002030302

Lange JR, Fabry B. 2013. Cell and tissue mechanics in cell migration. Experimental Cell Research 319:2418–2423. doi:10.1016/J.YEXCR.2013.04.023

Lawson CD, Ridley AJ. 2017. Rho GTPase signaling complexes in cell migration and invasion. Journal of Cell Biology 217:447–457. doi:10.1083/jcb.201612069

Lawson ND, Weinstein BM. 2002. In Vivo Imaging of Embryonic Vascular Development Using Transgenic Zebrafish. Developmental Biology 248:307–318. doi:10.1006/DBIO.2002.0711

Lee JSH, Hale CM, Panorchan P, Khatau SB, George JP, Tseng Y, Stewart CL, Hodzic D, Wirtz D. 2007. Nuclear Lamin A/C Deficiency Induces Defects in Cell Mechanics, Polarization, and Migration. Biophysical Journal 93:2542–2552. doi:10.1529/BIOPHYSJ.106.102426

Leitner L, Shaposhnikov D, Mengel A, Descot A, Julien S, Hoffmann R, Posern G. 2011. MAL/MRTF-A controls migration of non-invasive cells by upregulation of cytoskeleton-associated proteins. Journal of Cell Science 124:4318–4331. doi:10.1242/jcs.092791

Lin Y, Banno K, Gil C-H, Myslinski J, Hato T, Shelley WC, Gao H, Xuei X, Liu Y, Basile DP, Yoshimoto M, Prasain N, Tarnawsky SP, Adams RH, Naruse K, Yoshida J, Murphy MP, Horie K, Yoder MC. 2023. Origin, prospective identification, and function of circulating endothelial colony-forming cells in mice and humans. JCI Insight 8:e164781. doi:10.1172/jci.insight.164781

Mason DE, Collins JM, Dawahare JH, Nguyen TD, Lin Y, Voytik-Harbin SL, Zorlutuna P, Yoder MC, Boerckel JD. 2019. YAP and TAZ limit cytoskeletal and focal adhesion maturation to enable persistent cell motility. Journal of Cell Biology 218:1369–1389. doi:10.1083/jcb.201806065

Mathur J, Shenoy VB, Pathak A. 2020. Mechanical memory in cells emerges from mechanotransduction with transcriptional feedback and epigenetic plasticity. bioRxiv 2020.03.20.000802. doi:10.1101/2020.03.20.000802

Nakajima H, Yamamoto K, Agarwala S, Terai K, Fukui H, Fukuhara S, Ando K, Miyazaki T, Yokota Y, Schmelzer E, Belting H-G, Affolter M, Lecaudey V, Mochizuki N. 2017. Flow-Dependent Endothelial YAP Regulation Contributes to Vessel Maintenance. Developmental Cell 40:523–536.e6. doi:10.1016/J.DEVCEL.2017.02.019

Nasrollahi S, Walter C, Loza AJ, Schimizzi G V., Longmore GD, Pathak A. 2017. Past matrix stiffness primes epithelial cells and regulates their future collective migration through a mechanical memory. Biomaterials 146:146–155. doi:10.1016/j.biomaterials.2017.09.012

Neto F, Klaus-Bergmann A, Ong YT, Alt S, Vion A-C, Szymborska A, Carvalho JR, Hollfinger I, Bartels-Klein E, Franco CA, Potente M, Gerhardt H. 2018. YAP and TAZ regulate adherens junction dynamics and endothelial cell distribution during vascular development. eLife 7:e31037. doi:10.7554/eLife.31037

Neto F, Klaus-Bergmann A, Ong Y-T, Alt S, Vion A-C, Szymborska A, Carvalho JR, Hollfinger I, Bartels-Klein E, Franco CA, Potente M, Gerhardt H. 2017. YAP and TAZ regulate adherens junction dynamics and endothelial cell distribution during vascular development. bioRxiv 174185. doi:10.1101/174185

Nüsslein-Volhard C. 2006. Coming to life: how genes drive development. Kales Press.

Oakes PW, Wagner E, Brand CA, Probst D, Linke M, Schwarz US, Glotzer M, Gardel ML. 2017. Optogenetic control of RhoA reveals zyxin-mediated elasticity of stress fibres. Nature Communications 8:15817. doi:10.1038/ncomms15817

Ong YT, Andrade J, Armbruster M, Shi C, Castro M, Costa ASH, Sugino T, Eelen G, Zimmermann B, Wilhelm K, Lim J, Watanabe S, Guenther S, Schneider A, Zanconato F, Kaulich M, Pan D, Braun T, Gerhardt H, Efeyan A, Carmeliet P, Piccolo S, Grosso AR, Potente M. 2022. A YAP/TAZ-TEAD signalling module links endothelial nutrient acquisition to angiogenic growth. Nat Metab 4:672–682. doi:10.1038/s42255-022-00584-y

Pan D. 2010. The hippo signaling pathway in development and cancer. Developmental Cell 19:491–505. doi:10.1016/j.devcel.2010.09.011

Phillips JE, Santos M, Kanchwala M, Xing C, Pan D. 2022. Genome editing in the unicellular holozoan Capsaspora owczarzaki suggests a premetazoan function for the Hippo pathway in multicellular morphogenesis. bioRxiv 2021.11.15.468130. doi:10.1101/2021.11.15.468130

Phng L-K, Stanchi F, Gerhardt H. 2013. Filopodia are dispensable for endothelial tip cell guidance. Development 140:4031–4040. doi:10.1242/dev.097352

Pillay LM, Yano JJ, Davis AE, Butler MG, Ezeude MO, Park JS, Barnes KA, Reyes VL, Castranova D, Gore A V., Swift MR, Iben JR, Kenton MI, Stratman AN, Weinstein BM. 2022. In vivo dissection of Rhoa function in vascular development using zebrafish. Angiogenesis 1–24. doi:10.1007/s10456-022-09834-9

Porazinski S, Wang H, Asaoka Y, Behrndt M, Miyamoto T, Morita H, Hata S, Sasaki T, Krens SFG, Osada Y, Asaka S, Momoi A, Linton S, Miesfeld JB, Link BA, Senga T, Castillo-Morales A, Urrutia AO, Shimizu N, Nagase H, Matsuura S, Bagby S, Kondoh H, Nishina H, Heisenberg C-P, Furutani-Seiki M. 2015. YAP is essential for tissue tension to ensure vertebrate 3D body shape. Nature 521:217–221. doi:10.1038/nature14215

Posern G, Sotiropoulos A, Treisman R. 2002. Mutant actins demonstrate a role for unpolymerized actin in control of transcription by serum response factor. Molecular biology of the cell 13:4167–78. doi:10.1091/mbc.02-05-0068

Price CC, Mathur J, Boerckel JD, Pathak A, Shenoy VB. 2021. Dynamic self-reinforcement of gene expression determines acquisition of cellular mechanical memory. Biophysical Journal 120:5074–5089. doi:10.1016/J.BPJ.2021.10.006

Rapp BM, Saadatzedeh MR, Ofstein RH, Bhavsar JR, Tempel ZS, Moreno O, Morone P, Booth DA, Traktuev DO, Dalsing MC, Ingram DA, Yoder MC, March KL, Murphy MP. 2011. Resident Endothelial Progenitor Cells From Human Placenta Have Greater Vasculogenic Potential Than Circulating Endothelial Progenitor Cells From Umbilical Cord Blood. Cell Med 2:85–96. doi:10.3727/215517911X617888

Rosa A, Giese W, Meier K, Alt S, Klaus-Bergmann A, Edgar LT, Bartels-Klein E, Collins RT, Szymborska A, Coxam B, Bernabeu MO, Gerhardt H. 2022. WASp controls oriented migration of endothelial cells to achieve functional vascular patterning. Development 149. doi:10.1242/dev.200195

Ruehle MA, Eastburn EA, LaBelle SA, Krishnan L, Weiss JA, Boerckel JD, Wood LB, Guldberg RE, Willett NJ. 2020. Extracellular matrix compression temporally regulates microvascular angiogenesis. Science Advances 6:eabb6351. doi:10.1126/sciadv.abb6351

Sakabe M, Fan J, Odaka Y, Liu N, Hassan A, Duan X, Stump P, Byerly L, Donaldson M, Hao J, Fruttiger M, Lu QR, Zheng Y, Lang RA, Xin M. 2017. YAP/TAZ-CDC42 signaling regulates vascular tip cell migration. Proceedings of the National Academy of Sciences 114:10918– 10923. doi:10.1073/pnas.1704030114

Sivaraj KK, Dharmalingam B, Mohanakrishnan V, Jeong H-W, Kato K, Schröder S, Adams S, Koh GY, Adams RH. 2020. YAP1 and TAZ negatively control bone angiogenesis by limiting hypoxia-inducible factor signaling in endothelial cells. eLife 9:e50770. doi:10.7554/eLife.50770

Speight P, Kofler M, Szászi K, Kapus A. 2016. Context-dependent switch in chemo/mechanotransduction via multilevel crosstalk among cytoskeleton-regulated MRTF and TAZ and TGFβ-regulated Smad3. Nature Communications 7:11642. doi:10.1038/ncomms11642

Sunyer R, Conte V, Escribano J, Elosegui-Artola A, Labernadie A, Valon L, Navajas D, García-Aznar JM, Muñoz JJ, Roca-Cusachs P, Trepat X. 2016. Collective cell durotaxis emerges from long-range intercellular force transmission. Science 353:1157–1161. doi:10.1126/science.aaf7119

Sunyer R, Trepat X. 2020. Durotaxis. Current biology : CB 30:R383–R387. doi:10.1016/j.cub.2020.03.051

Totaro A, Panciera T, Piccolo S. 2018. YAP/TAZ upstream signals and downstream responses. Nat Cell Biol 20:888–899. doi:10.1038/s41556-018-0142-z

Valon L, Marín-Llauradó A, Wyatt T, Charras G, Trepat X. 2017. Optogenetic control of cellular forces and mechanotransduction. Nature Communications 8:14396. doi:10.1038/ncomms14396

van der Stoel M, Schimmel L, Nawaz K, van Stalborch A-M, de Haan A, Klaus-Bergmann A, Valent ET, Koenis DS, van Nieuw Amerongen GP, de Vries CJ, de Waard V, Gloerich M, van Buul JD, Huveneers S. 2020. DLC1 is a direct target of activated YAP/TAZ that drives collective migration and sprouting angiogenesis. Journal of cell science 133. doi:10.1242/jcs.239947

Wang K-C, Yeh Y-T, Nguyen P, Limqueco E, Lopez J, Thorossian S, Guan K-L, Li Y-SJ, Chien S. 2016. Flow-dependent YAP/TAZ activities regulate endothelial phenotypes and atherosclerosis. Proceedings of the National Academy of Sciences of the United States of America 113:11525–11530. doi:10.1073/pnas.1613121113

Wang X, Freire Valls A, Schermann G, Shen Y, Moya IM, Castro L, Urban S, Solecki GM, Winkler F, Riedemann L, Jain RK, Mazzone M, Schmidt T, Fischer T, Halder G, Ruiz de Almodóvar C. 2017. YAP/TAZ Orchestrate VEGF Signaling during Developmental Angiogenesis. Developmental cell 42:462–478.e7. doi:10.1016/j.devcel.2017.08.002

Webster KD, Ng WP, Fletcher DA. 2014. Tensional homeostasis in single fibroblasts. Biophysical Journal 107:146–155. doi:10.1016/j.bpj.2014.04.051

Weijts B, Gutierrez E, Saikin SK, Ablooglu AJ, Traver D, Groisman A, Tkachenko E. 2018. Blood flow-induced Notch activation and endothelial migration enable vascular remodeling in zebrafish embryos. Nature Communications 9:5314. doi:10.1038/s41467-018-07732-7

Yang C, Tibbitt MW, Basta L, Anseth KS. 2014a. Mechanical memory and dosing influence stem cell fate. Nature materials 13:645–52. doi:10.1038/nmat3889

Yang C, Tibbitt MW, Basta L, Anseth KS. 2014b. Mechanical memory and dosing influence stem cell fate. Nature materials 13:645–52. doi:10.1038/nmat3889

Yoder MC, Mead LE, Prater D, Krier TR, Mroueh KN, Li F, Krasich R, Temm CJ, Prchal JT, Ingram DA. 2006. Redefining endothelial progenitor cells via clonal analysis and hematopoietic stem/progenitor cell principals. Blood 109:1801–1809. doi:10.1182/blood-2006-08-043471

